# ENTPD3-specific CAR regulatory T cells for local immune control in T1D

**DOI:** 10.1101/2024.11.12.622951

**Authors:** Tom Pieper, Tobias Riet, Valerie Saetzler, Katharina Bergerhoff, Jenny McGovern, Louise Delsing, Mark Atkinson, Tracey Lodie, Lutz Jermutus, Fatih Noyan, Michael Hust, Irina Kusmartseva, Maike Hagedorn, Maren Lieber, Pierre Henschel, Viktor Glaser, Robert Geffers, Mingxing Yang, Julia Polansky-Biskup, Britta Eiz-Vesper, Agnes Bonifacius, Marc Martinez Llordella, Luke Henry, Daniela Penston, Artemis Gavriil, Thomas Grothier, Evanthia Nikolopoulou, Nikolaos Demertzis, Victoria Koullourou, Phillipa Cox, Bader Zarrouki, Marcella Sini, Janice Pfeiff, Isabelle Matthiesen, Matthias Hardtke-Wolenski, Elmar Jaeckel

## Abstract

Despite advances in Type 1 Diabetes (T1D) management such as hybrid closed loop systems, patients still face significant morbidity, reduced life expectancy, and impaired glucose regulation compared to healthy individuals or those with pancreas transplants.

Here we developed beta cell-specific Chimeric Antigen Receptors (CAR) targeting the antigen ectonucleoside triphosphate diphosphohydrolase 3 (ENTPD3) using a novel cell-based phage display methodology. ENTPD3 is highly expressed on beta cells of both early and progressed T1D patients. ENTPD3 CAR regulatory T cells (Tregs) homed, expanded and persisted in pancreatic islets in a T1D mouse model (NOD) and completely prevented disease progression. Human ENTPD3 CAR Tregs displayed a stable regulatory phenotype, strong activation, and suppression. Importantly, ENTPD3 CAR T cells recognised and were fully activated by human islets.

This approach holds great promise as a durable treatment option for patients with prediabetes, new-onset diabetes, or those undergoing beta cell replacement therapy.

## Introduction

Patients diagnosed with Type 1 Diabetes (T1D) continue to face substantial challenges, experiencing significant morbidity primarily attributable to the complications arising from fluctuating blood glucose levels and overall see a reduction in life expectancy by a decade^1-3^.

Although technological advancements, such as Continuous Glucose Monitoring (CGM) sensors and hybrid closed-loop pumps, have improved glycaemic control, particularly during nocturnal periods, they fail to replicate the robust metabolic control achieved by endogenous beta cells^4,5^.

While pancreas, islet or stem cell derived beta cell transplantation offers a potential biological replacement for beta cells, the associated challenges and risks, including immune rejection and the need for systemic immunosuppression, underscore the demand for innovative therapeutic approaches^6^.

Disease modifying therapies and immune modulation strategies, particularly in the context of new-onset T1D, have recently demonstrated success in clinical trials, albeit leading to only transient preservation of minimally stimulated c-peptide responses^7,8^.

Clinical trials administering polyclonal regulatory T cells (Tregs) in patients with T1D have shown a good safety profile but limited efficacy^9,10^. Pre-clinical data has shown superiority of antigen-specific Tregs over polyclonal Tregs in the NOD model of T1D and Tregs with beta cell-specific T cell receptors (TCR) have shown promise in establishing prolonged local tolerance in pre-clinical models^11-13^ but challenges arising from Major Histocompatibility Complex II (MHC II) polymorphism and the heterogeneity of immune responses in T1D patients have hindered the development of a viable clinical product for over two decades.

Recent advancements in the use of Chimeric Antigen Receptor (CAR) Treg cells have demonstrated success in achieving local immune control in murine and humanized mouse models. MHC I A*02 specific CAR-Tregs, activated independently of MHC II restriction, exhibit localized accumulation and prevented allograft rejection in mouse models and humanized mice^14-16^. Ongoing clinical trials exploring the use of A2-CAR-Tregs in kidney and liver transplantation provide a foundation for extending this approach to the context of T1D.

Herein, we identified ectonucleoside triphosphate diphosphohydrolase 3 (ENTPD3) as beta cell specific target and generated CARs directed against ENTPD3 using a novel phage display approach. We provide evidence for the antigen-specific functionality and safety of ENTDP3 CAR-Tregs *in vivo, in vitro* and *ex vivo*. Thus, ENTPD3-specific CAR Tregs offer a potential avenue for achieving durable, targeted and pancreas-specific immune control in patients with T1D.

## Results

### ENTPD3 as a Potential Target for Islet-Specific CAR-Treg Therapies

Previous attempts to develop CAR therapies targeting insulin, the most specific protein associated with beta cells, were ineffective in preventing or treating T1D^17^. In silico research suggested that ENTPD3, a surface protein, is highly expressed in both mouse and human beta cells^18-20^.

It is critical for potential therapeutic targets to be expressed at all stages of T1D pathophysiology. We demonstrated by histological analysis of pancreatic samples from patients at various stages of T1D, obtained through the nPOD (Network for Pancreatic Organ Donors with Diabetes) consortium, that ENTPD3 is consistently co-expressed with insulin across multiple stages of disease, from the autoantibody-positive stage to stage 3 (Fig. 1AB). Notably, some expression was detected in non-beta cells, during established, long duration T1D (Fig. 1A).

**Figure 1:**
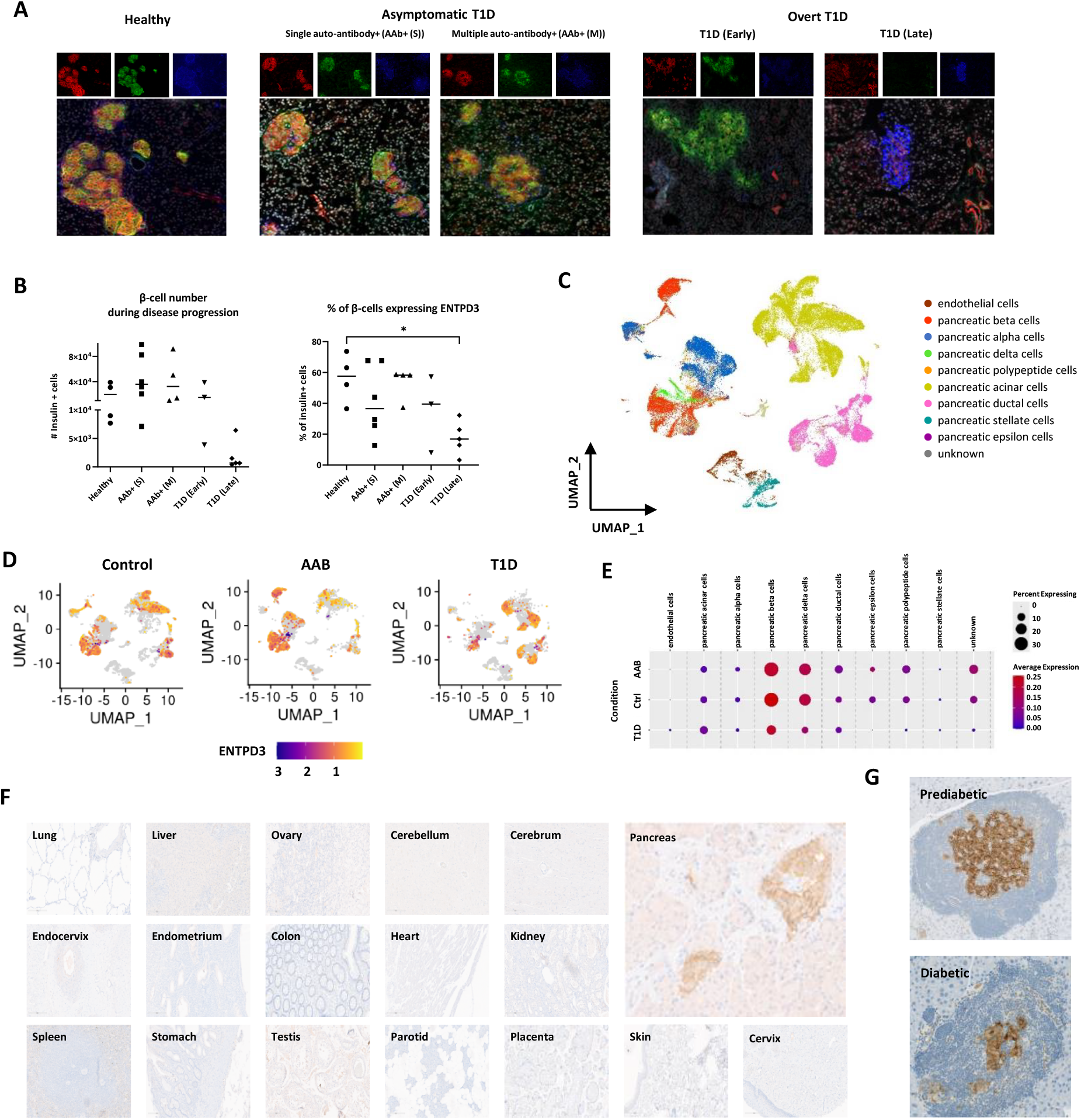
ENTPD3 as potential target for islets-specific CAR-Treg therapies. **(A)** Human pancreatic sections (nPOD) of individuals with different disease progression (Single Autoantibody positive: AAb+ (S); Multiple Autoantibody positive: AAb+ (M); Early and late T1D) were stained with anti-ENTPD3 antibody (red). Insulin (green) and glucagon (blue) were co-stained. Non-diabetic sample served as control. **(B)** Quantification of islets staining for ENTPD3. Left: Number of beta cells (Insulin+). Right: Percentage of beta cells expressing ENTPD3. n per condition = 3-6. **(C-E)** ENTPD3 expression of human pancreatic cells of nPOD samples. **(C)** UMAP visualization of cellular clusters of human pancreatic islets. **(D)** UMAP visualizations of ENTPD3 expression in islets cells of healthy (Control), autoantibody-positive (AAB) and T1D individuals (T1D). **(E)** Quantification of D. Dot size resembling percent of cells expressing ENTPD3 within cell cluster. Colours indicating average expression strength. **(F)** Tissue micro array (TMA) of different human organs stained for ENTPD3 (brown) show that membranous ENTPD3 expression is highly specific to pancreatic islets. Hematoxylin counterstain in blue. **(G)** Pancreatic sections of prediabetic and diabetic NOD mice were stained for ENTPD3 (brown) and counterstained with hematoxylin (blue).

The protein expression of ENTPD3 was corroborated by re-analysis of single-cell RNA sequencing data from patients at different stages of T1D (Fig. 1C-E)^21^, showing strong expression in beta cells across both autoantibody-positive and overt T1D stages as well as in delta cells as well. We confirmed specific expression of ENTPD3 protein in human pancreatic islets by immunohistochemistry analysis of a wide panel of human tissues (Fig. 1F). Finally, we confirmed that in prediabetic and diabetic NOD mice, ENTPD3 was expressed in islets, with minimal expression detected in surrounding ductal cells (Fig. 1G). These findings indicate that the NOD mouse model is suitable for proof-of-concept testing of ENTPD3-targeted CAR-Tregs.

### Generation of ENTPD3-Specific Single-Chain Variable Fragment (scFv) Binders via Protein-Cell Based Phage Display

Traditionally, CAR therapies have relied on a limited number of established monoclonal antibodies^22^. In contrast, our approach sought to generate diverse scFv binders and assess their functional properties. By employing human phage display libraries, we expedited the translation of binders into human CARs for clinical trials without requiring murine-to-human adaptation. Previous work from this lab demonstrated that successful panning against human peptide sequences can result in the selection of binders that cannot recognise intact proteins on cell surfaces ^23^. Consequently, we developed an assay that enriched scFv-phage binders against correctly folded surface proteins on target cells (Suppl. Fig. 1A) using cell sorting following rigorous pre-absorption and washing (Suppl. Fig. 2). This method produced a diverse array of ENTPD3-specific binders (Fig. 2A).

**Figure 2:**
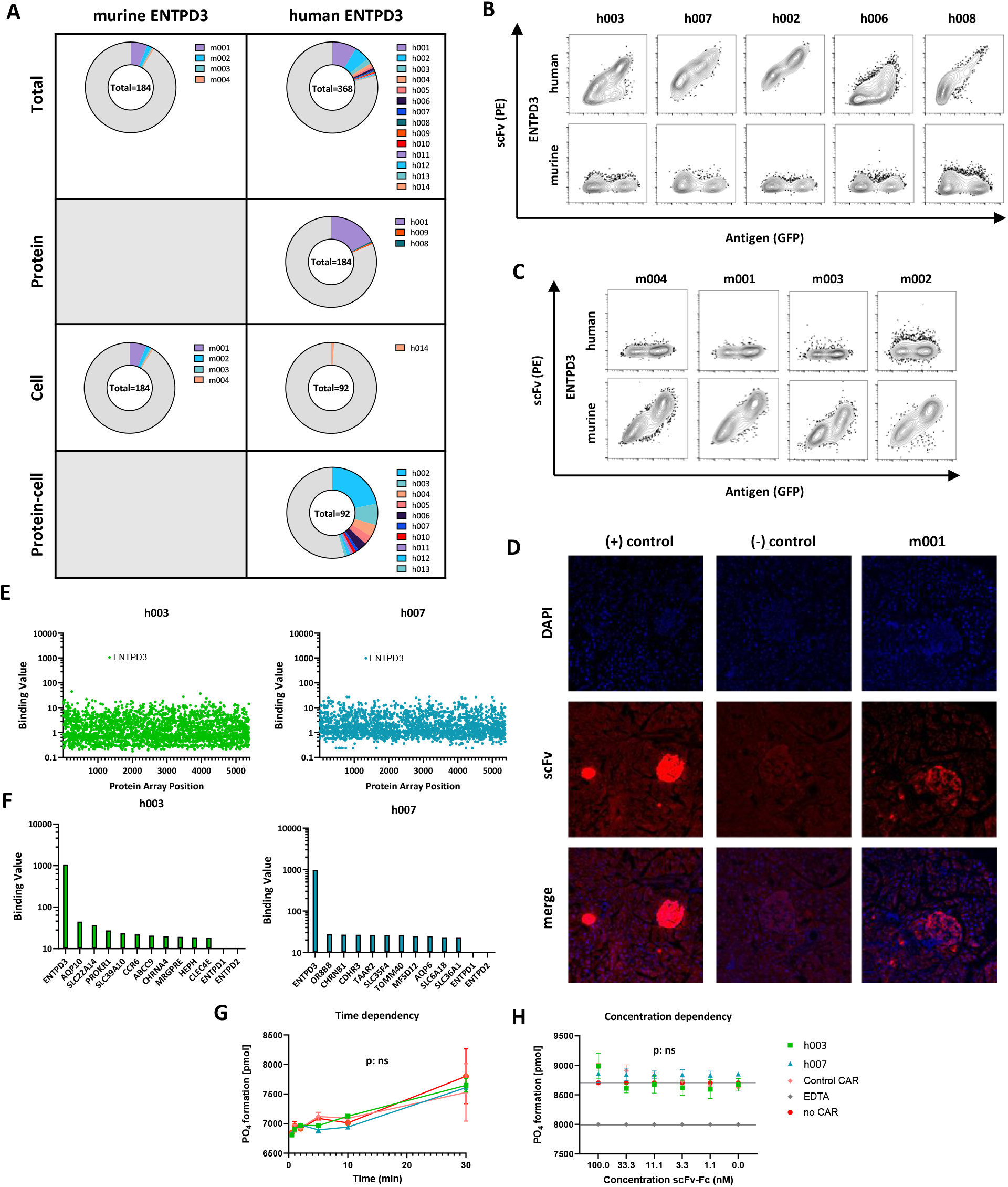
Generation of ENTPD3-specific scFv binders by protein- and cell-based phage display. **(A)** Outcome of scFv binders by different panning strategies for murine ENTPD3 and human ENTPD3. Total number of analysed scFv binders per strategy are given in the centre of each plot. Proportions of unique scFv clones (color-coded) and non- and unspecific binders (grey). **(B-C)** Binding of human **(B)** and murine **(C)** ENTPD3-specific scFv binders was measured by flow cytometry on human or murine ENTPD3 expressing HEK293T cells, respectively. ENTPD3 expression on HEK cells was reported by coexpression of eGFP. Representative plots of selected binders. **(D)** Staining of C57BL/6 pancreatic sections by murine ENTPD3-specific binder m001. Insulin-specific FITC-conjugated antibody was used as positive control (left). Incubation with scFv m001 was followed by FITC-conjugated secondary antibody (right). Staining only secondary antibody served as negative control (middle). **(E)** Membrane Proteome Array (MPA) screening was performed by testing of human ENTPD3 binders. Binding values are given for clones h003 and h007 in an scFv-Fc format for 5,372 distinct human membrane protein clones. **(F)** Binding values of MPA for ENTPD3 and 10 following proteins with highest values and ENTPD1 and ENTPD2. **(G-H)** ENTPD3 enzymatic activity in the presence of soluble ENTPD3, ADP and scFv-Fc of h003 or h007. ENTPD3 activity (PO_4_ formation) was measured by malachite green phosphate assay. Measured in triplicates. **(G)** Phosphate formation up top 30 min after incubation with 3.3 nM ENTPD3-specific scFv-Fc. No CAR and control CAR served as control. **(H)** Phosphate formation after incubation with varying concentrations (100 – 1 nM) of ENTPD3-specific scFv-Fc. Additionally, incubation with EDTA served as positive control for ENTPD3 inhibition. Data are presented as mean ± SD of triplicates. p values determined by two-way ANOVA of h003, h007 and Control CAR conditions, respectively. ns: P > 0.05.

We confirmed binding specificity using intact cells expressing ENTPD3 on the cell surface (Fig. 2BC). Furthermore, murine ENTPD3-specific binders were shown to stain islets of C57Bl/6 mice, confirming that selected binders could recognise ENTPD3 in the pancreas (Fig. 2D).

Interestingly, we were unable to identify binders that cross-reacted with human and murine ENTPD3 (Fig. 2BC), which is likely attributed to the low sequence homology between the human and murine extracellular domains (Suppl. Fig. 1B). Therefore, we continued with different scFvs for either murine-specific ENTPD3 to perform proof-of-concept experiments in NOD mice or human-specific ENTPD3 to further characterise of human CAR Tregs, respectively.

To rule out cross-reactivity with other membrane proteins, we screened the selected binders against 5,500 human membrane proteins using a membrane proteome array, confirming exclusive binding to ENTPD3 without cross-recognition of ENTPD1 or ENTPD2 (Fig. 2EF). ENTPD3 is an ectonucleotidase involved in extracellular nucleotide and ATP hydrolysis and thereby in purinergic signalling, contributing to the regulation of insulin secretion^19,24^. Consequently, it was essential to demonstrate that our scFv binders did not interfere with ENTPD3 enzymatic activity. We observed no evidence that scFv binding impaired ENTPD3 function, even when added at high concentrations (Fig. 2GH).

### Murine ENTPD3-Specific CARs are Activated upon Target Engagement and Prevent Cyclophosphamide-Induced Diabetes *In Vivo*

Murine ENTPD3-specific CARs were constructed using either CD8 hinge/transmembrane domains or IgG hinge/CD4 transmembrane domains, the latter featuring a mutated Fc receptor binding region to prevent nonspecific activation (Fig. 3AB). These second-generation CARs include a CD3zeta activation domain and a CD28 costimulatory domain, which are optimal for Treg activation, as demonstrated by Levings and colleagues^25^. CARs were tested in T cell hybridomas where NFAT activation induced eGFP expression via a minimal IL2 promoter (Suppl. Fig. 3A). CAR candidate m001 exhibited strong target-specific activation without background signalling in both hinge formats (Fig. 3CD) and was not stimulated by the human ENTPD1 (Suppl. Fig. 3B). m001 also recognised MIN6 beta cells (Suppl. Fig. 3CD) which were shown previously to express low levels of ENTPD3^19^. In murine CD4+ T cells, ENTPD3-specific CARs stimulated target-dependent proliferation and activation (Fig. 3E, Suppl. Fig. 4AB).

**Figure 3:**
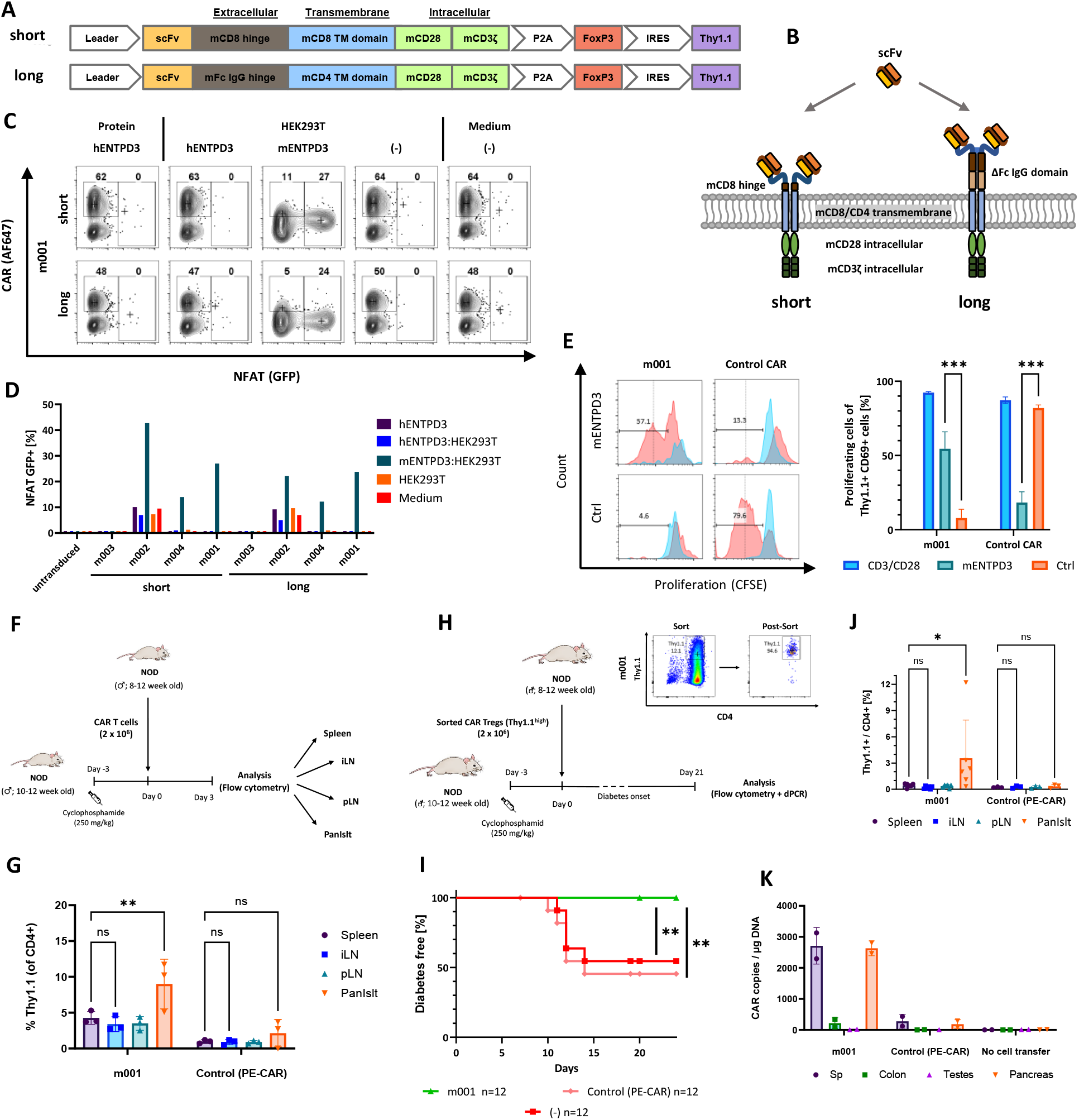
Murine ENTPD3-specific CAR are functionally stimulated upon target contact and prevent cyclophosphamide-induced diabetes in vivo. **(A)** Design of murine CD8-derived hinge CARs (short) and Fc-IgG-derived hinge CARs (long), containing a FOXP3 expression cassette separated by P2A cleavage side, and reporter gene Thy1.1 (CD90.1) under control of an IRES. **(B)** Illustration of both CAR constructs as expressed on the cell surface. **(C)** Representative plots of stimulation of m001 in short and long hinge CAR format, respectively, in NFAT-GFP reporter cell line. CAR stimulation shown as NFAT-controlled GFP expression and CAR expression shown as anti-Fab antibody AF647 staining. **(D)** Screening of further murine ENTPD-specific candidates for CAR stimulation in the aforementioned system. **(E)** Proliferation of CAR T cells measured as dilution of CFSE signal. CFSE labelled murine CD4+ CAR T cells (mL-m001 or Control (PE-specific) CAR) were stimulated on mENTPD3 or control antigen PE, or by aCD3/CD28 bead stimulus. Counts normalized to mode. Red: Stimulated CAR T cells (of CD69+ Thy1.1+). Blue: Unstimulated cells (of CD69-Thy1.1-). Left: Representative histograms. Right: Quantification of % Proliferation of stimulated CAR T cells (of CD69+ Thy1.1+). Mean ± SD, triplicates. p values determined by two-way ANOVA and multiple comparison testing (Tukey's test). (F) Schematic overview of setup for CAR T cell homing experiment in NOD mice. (G) Comparison of biodistribution of m001 and Control (PE-specific) CAR Teffs. CD4+ populations were analysed for percentage of Thy1.1+ cells. Data are presented as mean ± SD. n=3 per group. p values determined by two-way ANOVA and multiple comparison testing (Tukey's test). **(H)** Schematic overview of experimental setup for prevention of cyclophosphamide-induced diabetes in NOD mice. **(I)** Diabetes-free individuals over the course of the experiment. n=12 per group. Data from 8 independent experiments. p value determined by log-rank test for m001 CAR cTreg compared to Control (PE-specific) CAR cTreg and cyclophosphamide-only treated animals. **(J)** Comparison of biodistribution of m001 CAR-cTregs and Control (PE-specific) CAR cTregs at experimental endpoints (n = 3-6 per group). Data are presented as mean ± SD of Thy1.1+ in the CD4+ population. p values determined by two-way ANOVA and multiple comparison testing (Tukey's test). (K) Biodistribution of m001 CAR Tregs in ENTPD3-expressing tissues by digital PCR. Organs of two mice per group were saved at experimental endpoints and total gDNA was analysed by digital PCR. Copy numbers of CAR Treg-specific WPRE sequence per µg gDNA are displayed. P values for all experiments * P < 0.033, ** P < 0.002, *** P < 0.001.

**Figure 4:**
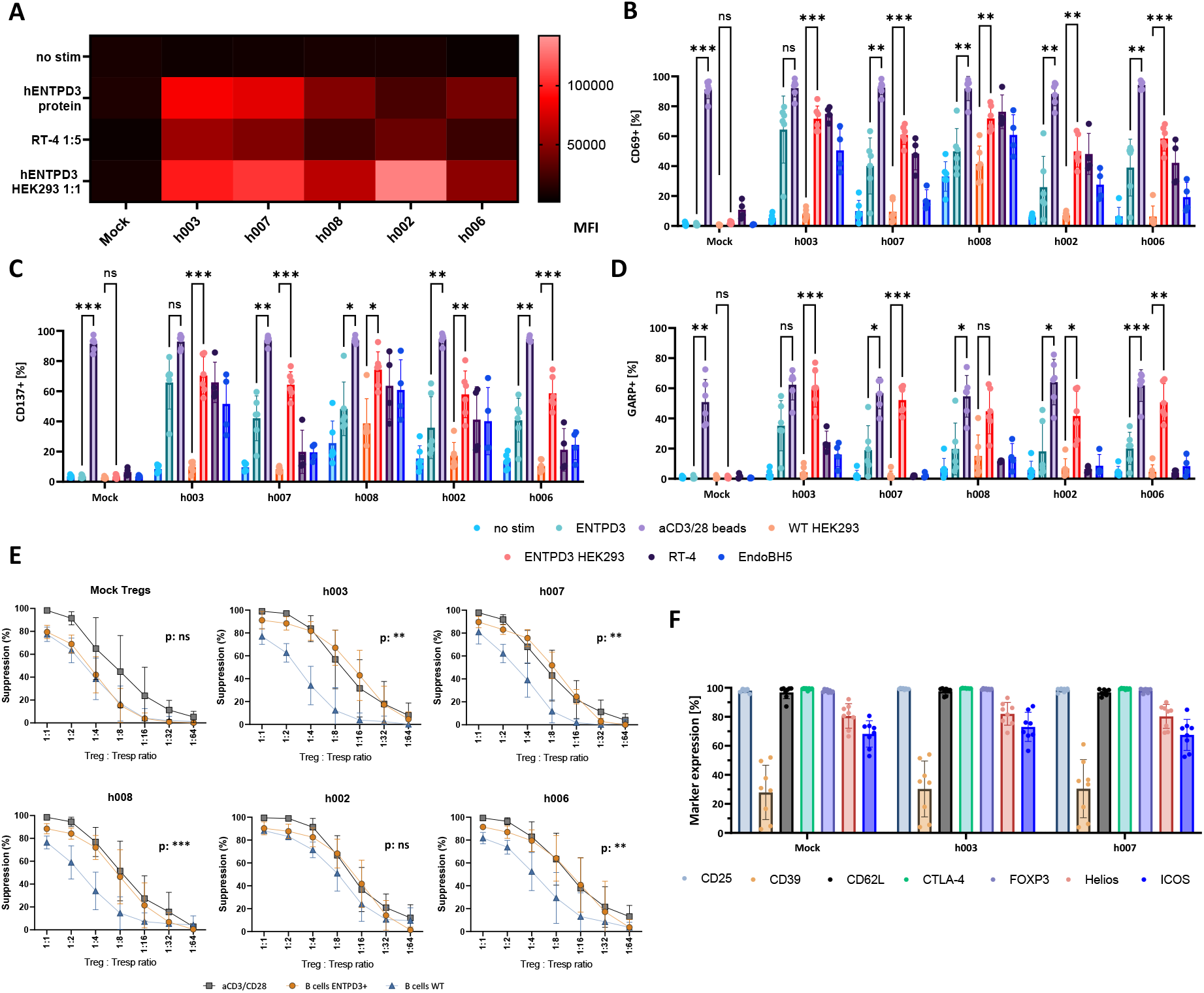
Human ENTPD3-specific CAR Tregs are functional and maintain Treg phenotype and suppressive function. **(A)** NFAT luciferase Jurkat cell line was transduced with CARs comprising different human ENTPD3-specific scFv (h003, h007, h008, h002 and h006). After 5 days cells were activated with human ENTPD3 extracellular domain peptide, cell lines expressing ENTPD3 (RT-4 and ENTPD3.HEK293T) at indicated ratios or left unstimulated. Activation of cells was determined by luminescence measurement. Heatmap shows relative activation levels in response to stimulus. Black represents the lowest levels of activation with red/pink showing highest levels of activation. Representative of 3 separate experiments. **(B-D)** Human ENTPD3 CAR Tregs were co-cultured with ENTPD3 extracellular domain, ENTPD3 expressing HEK293T cells, RT-4, EndoBH5 cells or controls (aCD3/aCD28 beads, WT HEK293T and no stimulation). After 24h activation markers CD69 **(B)**, CD137 **(C)** and GARP **(D)** were assessed by flow cytometry. Mean ± SD of n=6 donors. p values determined by two-way ANOVA and multiple comparison testing (Tukey's test). **(E)** CAR Tregs were co-cultured with WT B cell line, ENTPD3 expressing B cell line or aCD3/28 beads and decreasing numbers of CTV-labelled CD4+ Teffs for 5 days. Cells were assayed by flow cytometry and proliferation of Teffs was determined by CTV dilution. Graphs show percentage suppression by ENTPD3 CARs and mock transduced cells. Percentage suppression was calculated by normalizing proliferation of Teffs stimulated in the presence of Tregs to Teffs alone. Data are presented as mean ± SD of n=4-5 donors. p values determined by two-way ANOVA of B cells ENTPD3 and B cells WT conditions, respectively. **(F)** h003 and h007 CAR Tregs were collected and stained by flow cytometry for indicated markers. Percentage of each marker shown from the total CD4 population for Mock Tregs or gated on CAR+ cells for the CAR Tregs. Mean ± SD of n=8 donors. P values for all experiments * P < 0.033, ** P < 0.002, *** P < 0.001.

In a spontaneous diabetes NOD model, in which disease was synchronised by a single low-dose cyclophosphamide injection, ENTPD3-specific CAR-T cells (m001) showed strong early homing to pancreatic islets (day 3), unlike PE-specific control CAR-T cells (Fig. 3FG). In contrast, ENTPD3-specific CAR-T cells (m001) were absent in pancreatic lymph nodes. After 21 days, 15% of CD4+ cells in the pancreatic islets were ENTPD3 CAR-Tregs, reflecting a combination of homing, expansion, and persistence (Suppl. Fig. 4C).

In the cyclophosphamide-induced T1D model (Fig. 3H), m001 CAR-Tregs prevented diabetes onset, with no cases of T1D in the treatment group, compared to 50% incidence in control animals (Fig. 3I). In this experiment, we confirmed that ENTPD3 CAR-Tregs persisted long-term and were exclusively located in pancreatic islets (Fig. 3J). We observed no detectable homing to off-target tissues such as the testes or colon, as confirmed by digital PCR (Fig. 3K). Finally, murine ENTPD3 CAR-Tregs displayed a stable regulatory phenotype (Suppl. Fig. 5).

**Figure 5.**
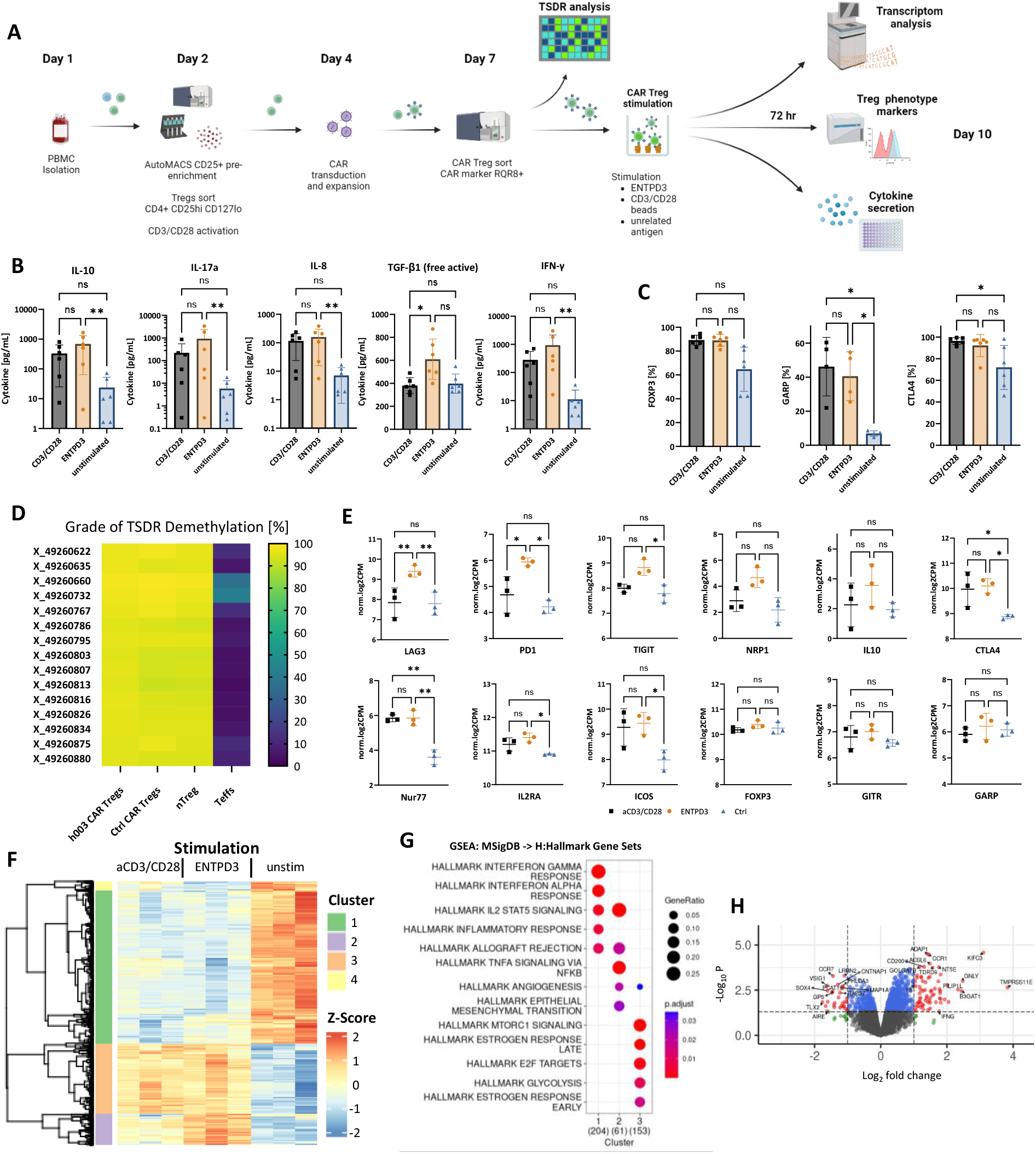
ENTPD3 CAR Tregs (candidate h003) maintain Treg-specific phenotype and gene expression upon stimulation. **(A)** Schematic overview and design of human CAR Treg stimulation assay containing TSDR, transcriptomics and phenotype markers and cytokine secretion readouts for testing of CAR h003. **(B)** Treg cytokine profile upon CAR and TCR-dependent stimulation. ENTPD3 CAR Treg cells were incubated with ENTPD3, PE (control antigen) or aCD3/CD28 beads. Cytokine concentration was determined by cytokine bead array. Data are presented as mean ± SD of n=6 donors. p values determined by two-way ANOVA and multiple comparison testing (Tukey's test). **(C)** Expression of Treg phenotype and activation markers FOXP3, GARP and CTLA4 upon stimulation measured by flow cytometry. ENTPD3-specific h003 CAR Tregs stimulated by aCD3/CD28 beads (black), ENTPD3 (orange) or unrelated control antigen (blue). Data presented as mean ± SD of 4-6 donors. p values determined by two-way ANOVA and multiple comparison testing (Tukey's test). **(D)** Analysis of demethylation status of Treg-specific demethylated regions (TSDR). Percentage of demethylation of different TSDRs is shown for ENTPD3 CAR Tregs (h003), Control (PE-specific) CAR Tregs, nTregs and Teffs. Mean of data from 3 donors. **(E)** Gene expression of Treg phenotype and activation genes upon stimulation of h003 CAR Tregs measured by RNA Sequencing. Black: TCR-dependent stimulation by aCD3/CD28 beads. Orange: CAR-specific stimulation by ENTPD3. Blue: Unrelated control antigen. Data mean ± SD of 3 donors. p values determined by two-way ANOVA and multiple comparison testing (Tukey's test). **(F)** Cluster analysis of transcriptome data. Z-score (color coded) shown for TCR-dependent stimulation by aCD3/CD28 beads, CAR-specific stimulation by ENTPD3 and unstimulated (unrelated control antigen). Data of 3 donors per condition. **(G)** Analysis of Human Molecular Signatures Database (MSigDB) Hallmark gene sets of major clusters 1, 2 and 3. Size of dots represent ratio of represented genes within each cluster. P value is color coded. **(H)** Volcano plot of differentially expressed genes in pairwise comparison of CAR-(ENTPD3) and TCR-(aCD3/CD28 beads) specific stimulation. p values for all experiments * P < 0.033, ** P < 0.002, *** P < 0.001.

### Generation of Human CARs Recognising ENTPD3

We deployed scFvs recognizing human ENTPD3 (Fig. 2B) to construct human CARs featuring either CD8 hinge and CD8 transmembrane region and performed functional characterization in Jurkat cells expressing luciferase under an NFAT-dependent minimal IL-2 promoter. Five CARs were identified that exhibited strong activation with no tonic signalling in the absence of antigen (Fig. 4A). Notably, CARs were also capable of recognising ENTPD3 on RT-4 cells, which endogenously express low levels of ENTPD3.

The five CARs identified in the screening stage were transduced into human nTregs. In activation assays using these cells, we observed robust upregulation of activation markers CD69, CD137, and the Treg-specific marker GARP following activation by ENTPD3 (Fig. 4B-D), with activation comparable to that of TCR stimulation. Coculture experiments of ENTPD3 expressing B cells, allospecific T effector cells (Teff) and ENTPD3 CAR Tregs demonstrated that ENTPD3 CAR Tregs could be activated by B cells expressing ENTPD3 to cross-suppress CD4+ Teffs activated by a TCR driven response to allogeneic B cells. This ability to control T cells with different specificities is a crucial function of CAR-Tregs given the need to suppress T cells reactivity against various beta cell antigens within the islets (Fig. 4E). Importantly, CAR transduction did not affect the stability of the regulatory phenotype, as evidenced by sustained expression of CD25, FOXP3, CTLA-4, Helios and ICOS as shown for CAR candidates h003 and h007 (Fig. 4F).

Second-generation CARs utilise partial activation motifs of the TCR complex, yet provide CD28 costimulatory signals. Therefore, we compared human ENTDP3-specific CAR Tregs in terms of CAR or TCR-based stimulation, focusing on the promising CAR candidate h003 more closely. CAR stimulated h003 CAR Tregs produced comparable levels of IL-10, IL-8, and IL-17a to those observed after TCR activation, with significantly increased TGF-β production (Fig. 5B). CAR Treg phenotype was stable after both CAR and TCR stimulation indicative of high FOXP3 and CTLA4 expression, and activation marker GARP was upregulated (Fig. 5C). Epigenetic demethylation of the TSDR region remained stable and indiscernible from untouched nTregs, further confirming phenotypic stability (Fig. 5D)^26^. We further investigated the molecular signalling pathways in CAR Tregs by RNA sequencing after CAR- and TCR-based activation. Interestingly, critical Treg-associated genes, such as CD25, GARP, ICOS, and CTLA4, displayed similar activation patterns, whereas LAG3, PD1 and TIGIT were more highly expressed after CAR activation (Fig. 5E). While most genes were similarly regulated following TCR and CAR stimulation, comparative cluster analysis identified genes that were uniquely upregulated following CAR stimulation in cluster 2 (Fig. 5F), characterised by genes involved in IL-2 signalling and NF-kB activation (Fig. 5G) and genes related to negative regulation of immune cells (Suppl. Fig. 6). Additionally, subtle differences in gene expression emerged as indicated by pairwise comparison (Fig. 5H).

**Figure 6:**
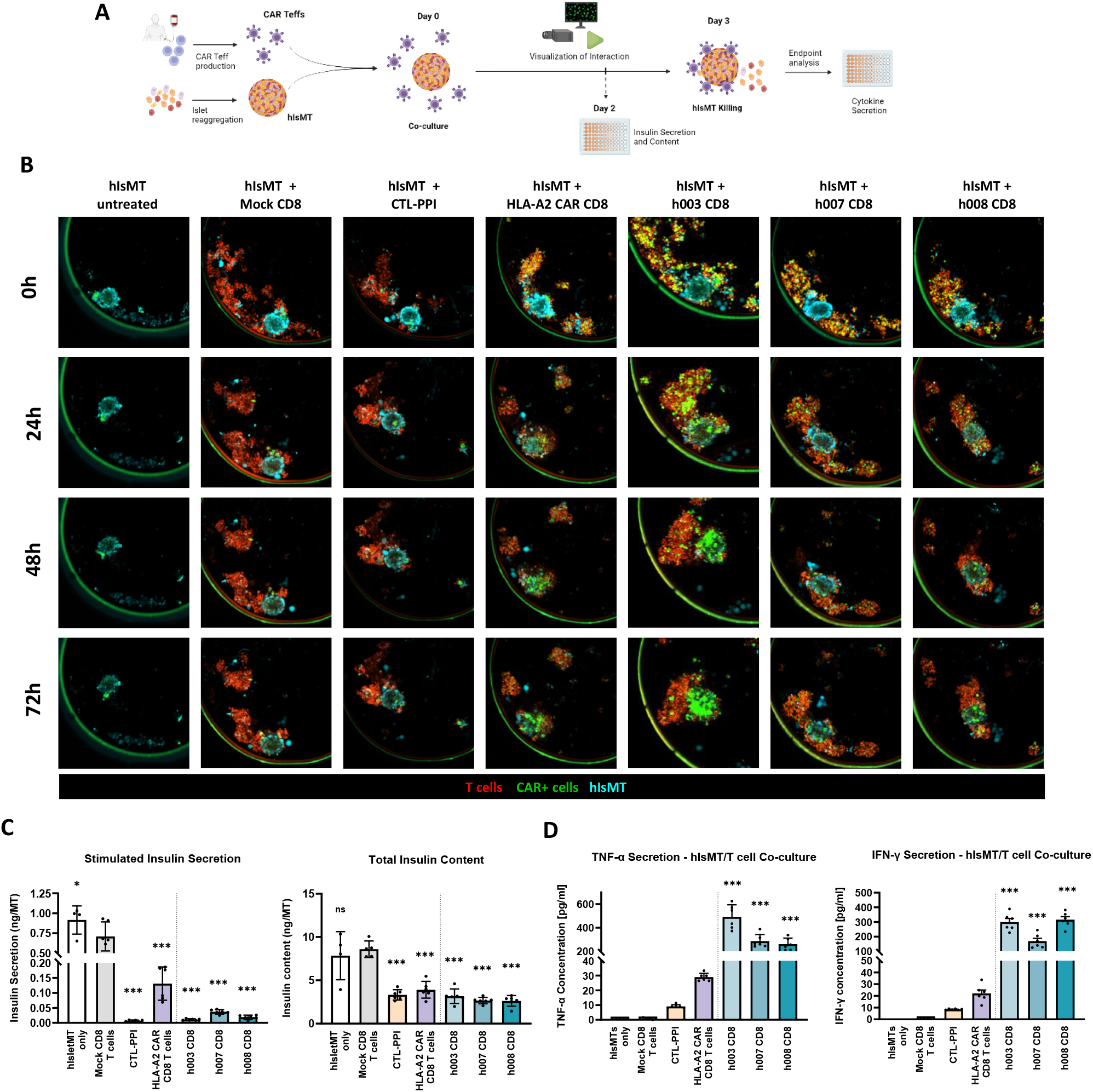
ENTPD3 CARs interact with human islets micro tissue (hIsMT) spheroids. **(A)** Schematic assay setup. **(B)** Human CD8+ T cells expressing either ENTPD3 CARs (h003, h007 or h008) or HLA-A2 CAR, and GFP reporter cassette were co-cultured with hIsMT spheroids for live imaging. Images show the hIsMT alone, with non-transduced CD8 T cells (Mock), preproinsulin (PPI) specific cytotoxic lymphocytes (CTLs) or CD8 T cells expressing HLA-A2 CAR or ENTPD3 CARs h003, h007 or h008 for timepoints 0, 24, 48 and 72 h. T cells are shown in red, CAR expression is shown in green and hIsMT are stained cyan. **(C)** After 48 h samples of co-cultures were lysed and stimulated insulin secretion (left) and total content (right) was determined by ELISA. Individual dots are technical replicates. Bars represent mean ± SD of n=4-6 **(D)** After 72 h supernatants were collected and assayed for IFN-γ (Ieft) and TNF-α (right). Individual dots are technical replicates. Bars represent mean ± SD of n=6. p values determined by One-way ANOVA with Dunnett’s multiple comparisons test comparing all conditions to mock CD8 T cells. Outliers were detected with ROUT’s outlier test (Q=5%). *p < 0.05, **p < 0.01, ***p < 0.001.

### *Ex vivo* Validation of Human ENTPD3 CAR Engagement and Activation

Based on these functional characteristics, three CAR candidates were selected for subsequent testing against human islets. Given the semi-quantitative nature of *in vitro* Treg function assays, we further characterised ENTPD3 CARs in Teffs, as quantitative differences in *in vitro* activation may correlate with variations in *in vivo* activity^27^. Notably, human CD4+ and CD8+ T cells transduced with the ENTPD3 CAR exhibited strong antigen-specific activation and proliferation (Suppl. Fig. 7)

As no relevant humanised T1D animal model exists, we were unable to test human ENTPD3 CAR Tregs *in vivo*. Instead, we assessed the capacity of ENTPD3 CAR-T cells to detect and interact with reaggregated human islets^28^ (Fig. 6A), comparing them to preproinsulin-specific CD8+ T cells^29^ and HLA-A*02-specific CAR T cells. ENTPD3 CAR T cells displayed time-dependent accumulation and invasion into the islets (Fig. 6B), leading to robust activation indicated by TNF-α and IFN-γ production and near-total destruction of beta cells and their function (Fig. 6CD). Remarkably, ENTPD3 CAR T cells exhibited stronger activation than did preproinsulin-specific CD8+ T cells and A2-specific CAR T cells. These experiments demonstrate that the amount of ENTPD3 on human beta cells and the signal transduction of ENTPD3 CARs are sufficient to activate T cells to full effector functionality.

## Discussion

Polyclonal regulatory T cell (Treg) therapies have demonstrated both safety and phenotypic stability in clinical trials targeting type 1 diabetes (T1D) and after organ transplantation^9,10,30^. Despite their excellent safety profiles, these therapies lack efficacy. It is estimated that just one in a million Treg cells will have specificity for a given MHC/self-peptide complex^31^.

In contrast, studies conducted over the past two decades have highlighted that Tregs recognizing beta cell antigens via their T cell receptors (TCRs), exhibit substantially higher potency than their polyclonal counterparts. Such antigen-specific Tregs have demonstrated long-lasting local immune regulation, successfully reversing even new-onset T1D in murine models^11-13^. Despite these promising results, translating antigen-specific Treg therapy into clinical practice has been hampered by the major histocompatibility complex class II (MHC II) restriction of TCRs, which significantly limits their applicability across diverse patient populations.

The advent of chimeric antigen receptors (CARs), which enable MHC-independent antigen recognition, has revolutionized immunotherapy in oncology^22,27^. The successful implementation of CAR T cells in oncology has paved the way for adapting this technology to Tregs to achieve local immune modulation in autoimmune diseases and transplantation. CAR Tregs, targeting mismatched HLA-A*02 molecules, have shown promise in inducing local immune control in transplantation models without systemic immunosuppression^14-16^. Encouragingly, these findings have led to clinical trials evaluating CAR Tregs in kidney and liver transplantation (LIBERATE NCT05234190).

In our study, ENTPD3 emerged as a highly promising target for beta cell-specific CAR Tregs. ENTPD3 is robustly expressed robustly in pancreatic beta cells and persists across all stages of T1D, from healthy individuals to patients with autoantibodies and recent-onset T1D. Importantly, ENTPD3 is also expressed in delta cells, even in cases of long-standing T1D, suggesting a broad yet specific expression pattern suitable for targeted immune regulation.

While current CAR approaches use binders derived from monoclonal antibodies, we employed the phage display technology to generate a large variety of scFv binders. While we and others have shown that scFv generated by phage display can be efficiently used in CAR studies^15,17,23,32,33^, these approaches deployed peptides and recombinant proteins which neglects the fact that most CARs target conformational epitopes on cell surface molecules. Therefore, we established a novel phage display approach directly on cells expressing ENTPD3, to generate a large variety of scFv binders recognising the correctly folded target on cells. This enabled us to choose the best binder for Treg activation and suppressive function. The use of human phage display libraries obviated the need for humanisation of CAR constructs, streamlining the translational process.

Off-target or on-target/off-tumor effects remain a significant concern in CAR T cell therapies, in immune oncology, where unintended tissue recognition can result in severe toxicities^34,35^. These toxicities are unlikely in Treg cell therapies; large doses of polyclonal Tregs have been shown to exhibit a strong safety profile, supporting the low risk associated with CAR Treg therapies^9,10,30^.Moreover, restricted expression of ENTPD3 in pancreatic islets, the absence of proinflammatory capabilities in Tregs and their dependency on local interleukin-2 (IL-2) signals at immune activation sites minimise the risk of adverse outcomes.

Comprehensive analyses, including tissue microarrays and membrane proteome profiling, confirmed that membranous ENTPD3 expression is restricted to the pancreas. Our investigations further demonstrated that ENTPD3 CARs do not inhibit the enzymatic activity of ENTPD3, mitigating potential unintended consequences of CAR binding on purinergic metabolism and signalling.

Although low ENTPD3 mRNA levels were detected in certain tissues, such as the testes and colon, these findings did not translate to protein expression, as confirmed by tissue microarray and in vivo biodistribution studies. ENTPD3 CAR Tregs predominantly localised to pancreatic islets and the spleen, with negligible signals detected in other tissues. Importantly, ENTPD3 CAR Tregs accounted for 15% of the CD4+ cells within pancreatic islets, underscoring their strong target tissue homing.

Mechanistically, ENTPD3 CAR Tregs differ from TCR Tregs in their mode of action. While TCR Tregs rely on interactions with MHC II and antigen-presenting cells, CAR Tregs act directly within the target tissue. This distinction accounts for the exclusive localisation of ENTPD3 CAR Tregs to the pancreatic islets and not to the draining lymph nodes. Our findings show that ENTPD3 CAR Tregs suppress immune responses through cross-suppression of CD4+ effector T cells with diverse antigen specificities and a critical feature in T1D where autoimmune responses target multiple beta cell antigens.

The concept of cross-suppression aligns with observations from transplantation models, where CAR Tregs activated by HLA-A*02 locally controlled immune responses against mismatched grafts with multiple allospecific T cells^15^. Moreover, a recent study by Levings et al. demonstrated that local activation of CAR Tregs can lead to the conversion of beta cell antigen-specific naïve T cells into Tregs^36^, a phenomenon previously described as “infectious tolerance” by Waldman et al^37^.

Transcriptomic analyses revealed that ENTPD3 CAR Tregs share activation profiles with TCR-activated Tregs, including upregulation of key regulatory genes such as IL-10, TGF-β, CTLA-4, and CD25. Notably, ENTPD3 CAR Tregs also exhibited increased expression of inhibitory receptors like LAG3, PD1, and TIGIT without signs of exhaustion^38^.

Concerns regarding the potential cytotoxicity of CAR Tregs^39,40^ were not observed in our study. ENTPD3-positive target cells, including HEK and B cells, did not exhibit signs of cytotoxicity following exposure to ENTPD3 CAR Tregs. This non-cytotoxic nature aligns with observations of ENTPD3 CARs^40^ and our in vivo data, where metabolic function remained stable despite high local densities of ENTPD3 CAR Tregs in NOD islets.

Ensuring phenotypic stability of human Tregs is critical for minimising the risk of differentiation into proinflammatory effector cells. By employing highly purified naïve Treg populations (CD25high CD127-CD45RA+), we mitigated risks associated with less stable Treg subsets^41,42^. ENTPD3 CAR Tregs exhibited stable demethylation in the Treg-specific demethylated region (TSDR), further demonstrating their phenotype stability^26^.

Future enhancements of phenotype stability could include incorporating FOXP3 transgenes into the CAR construct to bolster Treg stability and suppressor function under inflammatory conditions^43,44^ or strengthening the STAT5 signalling^45^.

Alternative beta cell-specific CAR targets, such as insulin and tetraspanin 7^17,23^, have been explored, but none have demonstrated comparable efficacy. ENTPD3 CAR Tregs offer a more versatile and non-MHC-restricted approach, paving the way for local immune control in T1D.

Our findings represent a significant advancement in the clinical translation of ENTPD3 CAR Tregs for T1D therapy. This approach could support endogenous beta cell regeneration in early T1D stages or complement transplantation of islets or stem-cell derived beta cells in advanced disease stages, potentially reducing reliance on systemic immunosuppression. ENTPD3 CAR Tregs hold promise as a “living drug” capable of achieving sustained, local immune regulation, representing a novel and transformative therapeutic option in the treatment of T1D.

## Acknowledgements

This work was supported by the ReSHAPE program, funded under the EU2020 program, with additional support from The Leona M. and Harry B. Helmsley Charitable Trust (2018PG-T1D06.3) and JDRF (3-SRA-2024-1529-S-B). Human CAR characterization was done in collaboration with Quell Therapeutics and Astra Zeneca. Quell Therapeutics supported the work at MHH as part of collaborative research support. Experiments with reaggregated human pseudoislets were done in collaboration with Quell Therapeutics and InSphero AG. This research was performed with the support of the Network for Pancreatic Organ Donors with Diabetes (nPOD), a collaborative type 1 diabetes research project funded by JDRF. Organ Procurement Organizations partnering with nPOD to provide research resources are listed at www.jdrfnpod.org/our-partners.php. The authors want to express their thank to Matthias Ballmaier of MHH’s Central Research Facility Cell Sorting for supervision and support. Maximilian Fuchs of Fraunhofer Institute for Toxicology and Experimental Medicine (ITEM) for providing help in RNAseq data interpretation and to the Hannover Biomedical Research School (HBRS) for attending the PhD project of Tom Pieper. The PhD thesis of Tom Pieper and Master thesis of Viktor Glaser contains parts of the data published here.

## References

1. Lind, M., et al. Glycemic control and excess mortality in type 1 diabetes. N Engl J Med 371, 1972–1982 (2014).

2. Beck, R.W., et al. The T1D Exchange clinic registry. J Clin Endocrinol Metab 97, 4383–4389 (2012).

3. Rawshani, A., et al. Mortality and Cardiovascular Disease in Type 1 and Type 2 Diabetes. N Engl J Med 376, 1407–1418 (2017).

4. Wadwa, R.P., et al. Trial of Hybrid Closed-Loop Control in Young Children with Type 1 Diabetes. N Engl J Med 388, 991–1001 (2023).

5. Choudhary, P., et al. Advanced hybrid closed loop therapy versus conventional treatment in adults with type 1 diabetes (ADAPT): a randomised controlled study. Lancet Diabetes Endocrinol 10, 720–731 (2022).

6. Reichman, T., et al. Glucose-dependent insulin production and insulin-independence in patients with type 1 diabetes infused with stem cell-derived, fully differentiated islet cells (VX-880). in EASD, Vol. 66 (Suppl1) S229 (Diabetologia, Hamburg, 2023).

7. Herold, K.C., et al. An Anti-CD3 Antibody, Teplizumab, in Relatives at Risk for Type 1 Diabetes. N Engl J Med 381, 603–613 (2019).

8. Ramos, E.L., et al. Teplizumab and beta-Cell Function in Newly Diagnosed Type 1 Diabetes. N Engl J Med 389, 2151–2161 (2023).

9. Bluestone, J.A., et al. Type 1 diabetes immunotherapy using polyclonal regulatory T cells. Science translational medicine 7, 315ra189 (2015).

10. Marek-Trzonkowska, N., et al. Administration of CD4+CD25highCD127-regulatory T cells preserves beta-cell function in type 1 diabetes in children. Diabetes Care 35, 1817–1820 (2012).

11. Tang, Q., et al. In Vitro-expanded Antigen-specific Regulatory T Cells Suppress Autoimmune Diabetes. J Exp Med 199, 1455–1465 (2004).

12. Tarbell, K.V., Yamazaki, S., Olson, K., Toy, P. & Steinman, R.M. CD25+ CD4+ T Cells, Expanded with Dendritic Cells Presenting a Single Autoantigenic Peptide, Suppress Autoimmune Diabetes. J Exp Med 199, 1467–1477 (2004).

13. Jaeckel, E., von Boehmer, H. & Manns, M.P. Antigen-Specific FoxP3-Transduced T-Cells Can Control Established Type 1 Diabetes. Diabetes 54, 306–310 (2005).

14. MacDonald, K.G., et al. Alloantigen-specific regulatory T cells generated with a chimeric antigen receptor. J Clin Invest 126, 1413–1424 (2016).

15. Noyan, F., et al. Prevention of Allograft Rejection by Use of Regulatory T Cells With an MHC-Specific Chimeric Antigen Receptor. Am J Transplant 17, 917–930 (2017).

16. Boardman, D.A., et al. Expression of a Chimeric Antigen Receptor Specific for Donor HLA Class I Enhances the Potency of Human Regulatory T Cells in Preventing Human Skin Transplant Rejection. Am J Transplant 17, 931–943 (2017).

17. Tenspolde, M., et al. Regulatory T cells engineered with a novel insulin-specific chimeric antigen receptor as a candidate immunotherapy for type 1 diabetes. J Autoimmun 103, 102289 (2019).

18. Docherty, F.M., et al. ENTPD3 Marks Mature Stem Cell-Derived beta-Cells Formed by Self-Aggregation In Vitro. Diabetes 70, 2554–2567 (2021).

19. Syed, S.K., et al. Ectonucleotidase NTPDase3 is abundant in pancreatic beta-cells and regulates glucose-induced insulin secretion. Am J Physiol Endocrinol Metab 305, E1319–1326 (2013).

20. Saunders, D.C., et al. Ectonucleoside Triphosphate Diphosphohydrolase-3 Antibody Targets Adult Human Pancreatic beta Cells for In Vitro and In Vivo Analysis. Cell Metab 29, 745–754 e744 (2019).

21. Fasolino, M., et al. Single-cell multi-omics analysis of human pancreatic islets reveals novel cellular states in type 1 diabetes. Nat Metab 4, 284–299 (2022).

22. Posey, A.D., Jr., Young, R.M. & June, C.H. Future perspectives on engineered T cells for cancer. Trends Cancer 10, 687–695 (2024).

23. Pieper, T., et al. Generation of Chimeric Antigen Receptors against Tetraspanin 7. Cells 12(2023).

24. Lavoie, E.G., et al. Identification of the ectonucleotidases expressed in mouse, rat, and human Langerhans islets: potential role of NTPDase3 in insulin secretion. Am J Physiol Endocrinol Metab 299, E647–656 (2010).

25. Dawson, N.A.J., et al. Functional effects of chimeric antigen receptor co-receptor signaling domains in human regulatory T cells. Science translational medicine 12(2020).

26. Polansky, J.K., et al. DNA methylation controls Foxp3 gene expression. Eur J Immunol 38, 1654–1663 (2008).

27. Hamieh, M., Mansilla-Soto, J., Riviere, I. & Sadelain, M. Programming CAR T Cell Tumor Recognition: Tuned Antigen Sensing and Logic Gating. Cancer discovery 13, 829–843 (2023).

28. Yesildag, B., et al. Liraglutide protects beta-cells in novel human islet spheroid models of type 1 diabetes. Clin Immunol 244, 109118 (2022).

29. Skowera, A., et al. beta-cell-specific CD8 T cell phenotype in type 1 diabetes reflects chronic autoantigen exposure. Diabetes 64, 916–925 (2015).

30. Sawitzki, B., et al. Regulatory cell therapy in kidney transplantation (The ONE Study): a harmonised design and analysis of seven non-randomised, single-arm, phase 1/2A trials. Lancet 395, 1627–1639 (2020).

31. Serr, I., et al. Type 1 diabetes vaccine candidates promote human Foxp3(+)Treg induction in humanized mice. Nat Commun 7, 10991 (2016).

32. De Paula Pohl, A., et al. Engineered regulatory T cells expressing myelin-specific chimeric antigen receptors suppress EAE progression. Cell Immunol 358, 104222 (2020).

33. Yoon, J., et al. FVIII-specific human chimeric antigen receptor T-regulatory cells suppress T- and B-cell responses to FVIII. Blood 129, 238–245 (2017).

34. Ghilardi, G., et al. T cell lymphoma and secondary primary malignancy risk after commercial CAR T cell therapy. Nat Med 30, 984–989 (2024).

35. Bonifant, C.L., Jackson, H.J., Brentjens, R.J. & Curran, K.J. Toxicity and management in CAR T-cell therapy. Mol Ther Oncolytics 3, 16011 (2016).

36. Wardell, C.M., et al. Short Report: CAR Tregs mediate linked suppression and infectious tolerance in islet transplantation. bioRxiv (2024).

37. Cobbold, S. & Waldmann, H. Infectious tolerance. Curr Opin Immunol 10, 518–524 (1998).

38. Lamarche, C., et al. Tonic-signaling chimeric antigen receptors drive human regulatory T cell exhaustion. Proc Natl Acad Sci U S A 120, e2219086120 (2023).

39. Boroughs, A.C., et al. Chimeric antigen receptor costimulation domains modulate human regulatory T cell function. JCI Insight 5(2019).

40. Wu, X., et al. CD39 delineates chimeric antigen receptor regulatory T cell subsets with distinct cytotoxic & regulatory functions against human islets. Frontiers in immunology 15, 1415102 (2024).

41. Liu, W., et al. CD127 expression inversely correlates with FoxP3 and suppressive function of human CD4+ T reg cells. J Exp Med 203, 1701–1711 (2006).

42. Sakaguchi, S., Vignali, D.A., Rudensky, A.Y., Niec, R.E. & Waldmann, H. The plasticity and stability of regulatory T cells. Nat Rev Immunol 13, 461–467 (2013).

43. McGovern, J., Holler, A., Thomas, S. & Stauss, H.J. Forced Fox-P3 expression can improve the safety and antigen-specific function of engineered regulatory T cells. J Autoimmun 132, 102888 (2022).

44. Henschel, P., et al. Supraphysiological FOXP3 expression in human CAR-Tregs results in improved stability, efficacy, and safety of CAR-Treg products for clinical application. J Autoimmun 138, 103057 (2023).

45. Kremer, J., et al. Membrane-bound IL-2 improves the expansion, survival, and phenotype of CAR Tregs and confers resistance to calcineurin inhibitors. Frontiers in immunology 13, 1005582 (2022).

