## Supplementary material for "ENTPD3-specific CAR regulatory T cells for local immune control in T1D": Methods and Supplemental Figures

##### Animals

NOD/ShiLtJ and C57BL/6J mice were bred and maintained under pathogen-free conditions in individual ventilated cages (IVC) at the central animal facility of Hannover Medical School, Hannover, Germany. All animals were bred from founders obtained from Jackson Laboratory (Bar Harbor, ME, USA). All animals were scored and monitored for general health, behaviour, body weight and specific experimental criteria. Blood glucose was measured by puncturing the tip of the tail and transferring a minor amount of blood to a Beurer blood glucose meter. Diabetic state was defined as persistent blood glucose levels ≥250 mg/dL. All mouse strains expressed Thy1.2 (CD90.2). Animal care, treatment and experimental procedures were performed in accordance with institutional, state, and federal guidelines.

##### Ethical statement

Human specimens were obtained from different healthy donors by MHH´s department of Transfusion Medicine and Transplant Engineering. Local ethical committee approval was received for this study. Informed consent was obtained from all participating subjects.

##### Immunofluorescence and Immunohistochemistry

nPOD samples were fixed in acetone for 10 minutes before air drying. Slides were rehydrated by washing with PBS and blocked with 5% normal goat serum. Slides were incubated with primary antibody cocktail (1:500 anti-ENTPD3 (Mouse) + 1:12,000 anti-glucagon antibody (92517, Abcam) either 1:50 Millipore anti-somatostatin antibody (clone YC7, Sigma-Aldrich) or Insulin antibody (IR002, Dako) in 1X PBS + 1% BSA for 1 hour at room temperature. Fluorescent secondary antibodies AF555 goat anti-Rabbit, AF647 goat anti-Mouse and AF488 goat anti-Rat or AF488 goat anti- Guinea Pig were prepared at 1:500 in 2% normal goat serum and incubated at room temperature for 45 minutes. After a wash step, sections were fixed for 5 min in 2% Paraformaldehyde (PFA). Sections were mounted in ProLong™ Diamond Antifade Mountant with DAPI (Invitrogen).

Mouse pancreas tissues from non-diabetic, early, and late diabetic mice were embedded in paraffin wax. IHC staining was performed using Ventana automated system (Discovery Ultra S/N 313108, Roche). Paraffin sections were de-paraffinized and rehydrated with stepwise decreasing concentrations of ethanol. Subsequently, antigen retrieval was performed using Cell Conditioning Solution (Roche) for 30 min at 95 °C. The tissues were then blocked with 1% BSA and endogenous peroxidase activity was quenched with Inhibitor CM (Roche) for 12 min. The slides were stained with anti-ENTPD3 antibody (AF4464, R&D systems) followed by secondary rabbit anti-sheep antibody (Ab6704, Abcam). Finally, HRP-conjugated anti-rabbit antibody (760-4311, Roche) was added followed by 3, 3′-diaminobenzidine (DAB) for signal development. The tissues were then counterstained with haematoxylin, mounted and visualized under light microscope.

Binding of scFv to pancreatic tissues was analysed by staining of cryosections of pancreatic tissues derived from C57Bl/6 mice. Tissue was incubated for 4 hr in 4% paraformaldehyde solution, followed by incubation in 30% sucrose solution overnight. Tissues were embedded in Tissue-Tek® O.C.T. Compound (Sakura Finetek) and snap frozen in liquid nitrogen. 5 µm sections were dissected using cryotome sectioning and mounted on glass slides. For staining sections were allowed to dry at room temperature 1 hr prior staining. They were subsequentially incubated with TBS + Tween20, 100mM glycine buffer and TBS + Tween20 containing 20% BSA, respectively. Slides were stained with 300 µL of soluble scFv supernatant for 1 hr followed by 300 µL of rabbit α-Myc tag antibody diluted 1:200 in TBS + Tween20 containing 20% BSA and with 300 µL of FITC-labelled goat α-rabbit antibody (Invitrogen) diluted 1:300 in TBS + Tween20 containing 20% BSA, respectively. Staining steps were separated by washing with TBS + Tween20. The slides were mounted with 1-2 drops of Fluoromount-G™ Mounting Medium, with DAPI (Thermo Fisher Scientific) and cover-slipped with a glass coverslip and analysed the next day with Axio Imager M1 fluorescence microscope (Zeiss).

**Analysis of Human Pancreas Analysis Program (HPAP) transcriptomic data**

For analysis of ENTPD3 expression in patients of different Type 1 Diabetes severity, single-cell RNA sequencing data obtained by *Fasolino et al*[1] from Human Pancreas Analysis Program (HPAP) samples of autoantibody-positive (AAB), T1D patients (T1D) and healthy individuals (Control) were analysed for ENTPD3 transcripts. UMAP visualization distinguished different cell types by cell-type classifier Garnett[2] as described by *Fasolino et al*[1]. ENTPD3 expression was either visualized per cell for different HPAP sample groups or as average expression.

**Human Tissue Micro Array**

RNAscope prequalified normal (non-malignant) 2 mm core diameter human tissue micro array (TMA) slides were purchased from Tristar Technology Group. Immunohistochemistry methods and protocols were set up on an automated Ventana Discovery Ultra autostainer (V12.5.4, Roche). Immunohistochemistry for detection of desired epitope was carried out according to the manufacturer's recommendation and all reagents except antibodies were Ventana products (Roche). Antigen retrieval was done before primary antibody hENTPD3 was added, followed by antibody block and anti-sheep linker and secondary anti-rabbit reagent, and DAB chromogenic detection in single staining. The slides were digitized using a Aperio XT whole slide scanner and Aperio ImageScope software (V12.3.3.5048, Technology Group).

**Phage display for generation of ENTPD3-specific scFv**

Cell- and protein-based phage display panning approaches were utilized to generate both murine and human ENTPD3-specific scFv binders for CAR constructs. 5 x $10^{11}$ phage particles of fully human phage library HAL9/10[3] were blocked in 150 µL panning block solution (PBS + 1% skim milk powder + 1% bovine serum albumin (BSA)) for 30 min. For protein panning blocked phage solution was transferred to ELISA plate wells precoated with extracellular domain (Gln 44-Pro 485) of human ENTPD3 protein (Sino Biological) and incubated for 1 h at room temperature (RT). ELISA plate was washed up to 30 times with PBS containing 0,1% Tween20 (PBS-T) and eluted with 150 µL of trypsin solution for 30 min at 37 °C. Cell panning was performed by using human or murine ENTPD3 expressing HEK293T cells. For depletion of unspecific scFv-phage particles, blocked phage solution was incubated with 2 x $10^7$ untransfected HEK293T cells 1 h at 4 °C on a rotator. Depleted phage solution was incubated with 2 x $10^7$ HEK293T cells transfected with target vectors containing either murine or humane ENTPD3-linked to eGFP by P2A cleavage site indicating target expression by eGFP coexpression (Suppl. Fig. 1CD) for 1 h at 4 °C on a rotator. After testing different washing strategies (Supplementary Fig. 2B) target expressing HEK293T cells were washed in PBS and sorted for GFP+ population by flow cytometry cell sorting (Suppl. Fig. 2C). Phage particles were eluted by adding 250 µL trypsin solution in PBS and incubation for 30 min at RT on a rotator. Eluates were either used for amplification of scFv-phage particles for further panning rounds or isolation of single scFv binders. Enrichment of scFv-phage pools was checked by flow cytometry (Supplementary Fig. 1A). Detailed cell panning procedure and amplification of enriched scFv-phage particles were described previously[4].

**Screening of ENTPD3-specific scFv binders**

Phage eluate was serially diluted, mixed with 50 µL XL1-blue MRF’ culture in exponential phase and incubated for 30 min at 37 °C without shaking. The mix was plated onto 2x YT A agar plates and incubated overnight at 37 °C. The next day single colonies were picked to inoculate 96-well U-bottom plate wells containing 150 µL of 2x YT medium containing glucose and ampicillin (2x YT-GA). Uninfected wells and wells inoculated with specific binder served as negative and positive control, respectively. The plates were sealed with breathable foil and incubated overnight at 37 °C with shaking at 1.4 x g. 10 µL was used to inoculate a new 96-well U-bottom plate filled with pre-warmed 150 µL 2x YT GA and incubated for 2 h at 37 °C with rotation at 1.4 x g. The plate was then centrifuged at 3000 x g for 10 min and the supernatant was removed. Pellets were resuspended in 180 µL of pre-warmed 2x YT containing 50 µM IPTG and ampicillin and incubated overnight at 30 °C with rotation at 1.4 x g to produce soluble scFv.

HEK293T cells transfected with target vectors containing either murine or humane ENTPD3-linked to eGFP by P2A cleavage site were harvested and mixed with untransfected HEK293T cells to obtain a cell mixture containing 50% target positive cells. 5 x $10^4$ cells were transferred to each well of a 96-well V-bottom plate and resuspended in 60 µL soluble scFvs of the production plate and incubated for 20 min at RT. The plates were washed with MACS buffer and then stained with 1:200 PE-labelled α-His-Tag secondary antibody (GG11-8F3.5.1, Miltenyi Biotec) binding the His-Tag of scFv and analysed by flow cytometry.

For analysis of scFv-phage pools, 60 µL of phage amplificate was used instead of soluble scFv, and stained by 1:100 PE-labelled α-M13 major coat protein antibody (RL-ph1, Santa Cruz Biotechnology).

**Membrane proteome array**

Membrane Proteome Array (MPA) screening was conducted at Integral Molecular, Inc. to check for specific binding of solubilized scFv-Fcs of ENTPD3-binders h003 and h007. The MPA is a protein library composed of 5,372 distinct human membrane protein clones, each overexpressed in live cells from expression plasmids. Each clone was individually transfected in separate wells of a 384-well plate followed by a 36h incubation[5]. Cells expressing each individual MPA protein clone were arrayed in duplicate in a matrix format for high-throughput screening. Before screening on the MPA, the concentration of scFv-Fc for screening was determined on cells expressing positive (membrane-tethered Protein A) and negative (mock-transfected) binding controls, followed by detection by flow cytometry using a fluorescently-labelled secondary antibody. Each test ligand was added to the MPA at the predetermined concentration, and binding across the protein library was measured on an Intellicyt iQue using a fluorescently labelled secondary antibody. Each array plate contains both positive (Fc-binding) and negative (empty vector) controls to ensure plate-by-plate reproducibility. Test ligand interactions with any targets identified by MPA screening were confirmed in a second flow cytometry experiment using serial dilutions of the test antibody, and the target identity was re-verified by sequencing.

**ENTPD3 enzymatic activity assay**

ENTPD3 enzymatic activity in the presence of solubilized scFv-Fc for h003, h007 and an irrelevant scFv-Fc binding the HIV envelope glycoprotein GP120 (control CAR) was evaluated by measuring the release of phosphate which occurs when ENTPD3 converts ADP to AMP. scFv-Fc were incubated with soluble ENTPD3 (R&D Systems) and ATP and the released phosphate was measured using the Malachite green phosphate assay kit (Sigma-Aldrich).

Two experiments were performed to evaluate the effects of the FC-scFv on ENTPD3. In the first experiment, ENTPD3 (R&D Systems) diluted in 25 mM Tris, pH 7.4, 5 mM CaCl2 containing ADP (25 mM) was incubated in a concentration range of 0 to 100 µM Fc-scFv in a 96 well plate. An EDTA control (25 µM) (Invitrogen) was added as reference for complete ENTPD3 inhibition. Malachite green solution was added to the plate at room temperature and absorbance was measured at 620 nm and 800 nm on a plate reader, SpectraMax i3 (Molecular Devices) after 30 min incubation. The amount of released phosphate was calculated for the concentration range of FC-scFvs in GraphPad Prism 10 (GraphPad Software Inc.)

In the second experiment ENTPD3 enzymatic activity was measured over time in presence of the scFvs-Fc. ENTPD3 (R&D Systems) diluted in 25 mM Tris, pH 7.4, 5 mM CaCl2 containing ADP (25 mM), Fc-scFv (3,3 nM) and Malachite green solution were added in 96 well plate at room temperature. Absorbance was measured at 620 nm and 800 nm on a plate reader SpectraMax i3 (Molecular Devices) after 30 sec, 1 min, 2 min, 5 min, 10 min and 30 min.

**CAR design**

ENTPD3-specific binders were either cloned in human or murine second-generation CAR constructs containing human or murine CD3- and CD28-derived signalling domains and a short or long hinge domain, respectively. Long hinge domain was constituted by human or murine IgG-Fc domains while short hinge domains were constituted out of CD8-derived (murine CAR) or CD4-derived (human CAR) hinges. Murine CAR constructs for generation of murine CAR Tregs had additional FOXP3 gene to lock Treg phenotype separated by P2A cleavage site. Human CAR constructs either contained additional FOXP3 or did not, based on the experiment. Reporter genes Thy1.1 (CD90.1) and RQR8 (CD20/CD34) preceded by internal ribosomal entry site (IRES) were used for murine and human CAR constructs, respectively. Gamma-retroviral particles for transduction of CARs into target cells were produced as described before[6].

**CAR functionality assay by NFAT-GFP reporter cell line**

CARs were checked for correct target-dependent stimulation in murine T cell-derived hybridoma cell lines which contain an artificial gene expression cassette with GFP expressed under the control of NFAT binding sites and a minimal IL-2 promoter and therefore report CAR stimulation by GFP expression[7]. Hybridoma cells were transduced by spin transduction as described earlier[8] and incubated for 24 h in wells coated with either extracellular domain (Gln 44-Pro 485) of human or murine ENTPD3 protein (Sino Biological) or HEK293 cells expressing murine or human ENTPD3 (Suppl. Fig. 1EF), murine beta cell line MIN6 or left untreated, respectively. Cells were harvested and stained with HER2 antibody (PE, 24D2, Biolegend) for off-gating of HER2 expressing HEK293T cells and αFab antibody (AF647, polyclonal, Jackson) for staining of extracellular CAR domain on hybridoma cells. Cells were analysed for CAR and endogenous GFP expression by flow cytometry.

**ENTPD3-specific CAR T cells in NOD mice**

For evaluation of CAR activation in murine T cells CD4+ and CD8+ Teffs were purified from murine spleenocytes using an MojoSort™ Mouse CD3 T Cell Isolation Kit (Biolegend). Cells were activated with aCD3/CD28 bead-based murine Treg expansion kit (Miltenyi) and transduced with retroviral CAR vectors. 48 h after transduction, CAR T cells were debeaded from αCD3/αCD28 activation beads. CAR Teffs were harvested and stained with CFSE for study of proliferation. 5 x 104 stained cells were seeded per well into a 96-well plate that had previously been coated with 1 µg/mL recombinant protein in 200 µL PBS for 2 h at 37 °C. As a positive control, the cells were mixed 1:1 with aCD3/CD28 Dynabeads in a separate uncoated well. Cells were incubated for 72 h at 37 °C. The cells were then stained for live cells (Fixable viability dye eFluor 780, Invitrogen), CD4 (BV421, L3T4, Biolegend), CD8 (PE, 53-6.7, BD Biosciences), Thy1.1 (AF647, OX-7, Biolegend) and CD69 (PE/Cy7, H1.2F3, Biolegend) and cell activation and proliferation were determined by flow cytometry.
Homing capabilities of murine CARs were investigated using CAR Teffs first. CD4+ T cells were isolated from NOD/ShiLtJ splenocytes using the MojoSort™ Mouse CD4 T Cell Isolation Kit

(Biolegend). Cells were activated with aCD3/CD28 bead-based murine Treg expansion kit (Miltenyi) for 48 h and transduced with retroviral particles of murine CAR constructs lacking the additional FOXP3 via spin inoculation (850× g, 32 °C, 1.5 h, MOI = 25) using protamine sulfate (4 µg/mL, Sigma-Aldrich). 24 h after transduction cells were debeaded, washed and injected intravenously into male NOD mice treated intraperitoneal with Cyclophosphamide (Sigma Aldrich) in a concentration of 250 µg per g body weight 72 h in advance. After 72 h mice were sacrificed, lymphocytes were isolated from spleen (Sp), inguinale lymph nodes (iLN), pancreatic lymph nodes (pLN) and pancreatic islets (PanIslt), stained for live cells (Fixable viability dye eFluor 780, Invitrogen), CD4 (BV421, L3T4, Biolegend), CD8 (FITC, 53-6.7, Biolegend), CD62L (BV510, MEL-14, Biolegend) and Thy1.1 (PE, OX-7, Biolegend) and analysed by flow cytometry. For isolation of lymphocytes from pancreatic islets the pancreas was perfused with collagenase solution () and incubated at 37 °C up to 10 min with short disruptive shaking every 2 minutes. Digestion was stopped by adding 25 mL ice-cold PBS and washed 2 times by centrifugation at 300 x g for 30 seconds and resuspension in ice-cold PBS. After washing, the resuspended suspension was filtered through a 70 µm filter. The filter was inverted and the islets were eluted with RPMI medium into a 10 cm Petri dish and islets were picked under microscope. Islets were disrupted in non-enzymatic cell dissociation solution (Sigma Aldrich) and incubated on a rocker at 37°C for 10 min. The dissociated cells were washed 2 times in RPMI medium and analysed analogously to lymphocytes isolated from lymphatic organs. For therapeutic experiments CD4 T cells of NOD mice were isolated as described above and transduced with CAR constructs containing ectopic FOXP3 gene to generate converted Tregs (CAR cTregs). 72 h after transduction CAR cTregs were stained by antibodies and CD4 (AF488, RM4-5, Biolegend) and Thy1.1 (AF647, OX-7, Biolegend) population was sorted using FACS Fusion cell sorter (BD Biosciences) and cells were transferred to the animals treated as described above. At experimental end points lymphocytes were collected from organs as described above, stained for live cells (Fixable viability dye eFluor 780, Invitrogen), CD4 (BV421, L3T4, Biolegend), CD8 (FITC, 53-6.7, Biolegend), FOXP3 (AF647, MF-14, Biolegend) and Thy1.1 (PE, OX-7, Biolegend) and analysed by flow cytometry. Phenotype of murine CAR cTregs converted by ectopic FOXP3 expression was evaluated previously by antibody staining of live cells (Fixable viability dye eFluor 780, Invitrogen), Thy1.1, FOXP3, CD25, CD62L, CD69, CTLA4 and GITR (all Biolegend) and subsequent analysis by flow cytometry.

**Digital PCR for biodistribution analysis**

Tissues (pancreas, colon, spleen and testes) from mice treated with ENTPD3 CAR-cTregs, Control (PE-CAR)-cTregs or no cell transfer, were homogenised using the Tissuelyser II (QIAGEN) in RLT Plus buffer (QIAGEN). They were then processed for DNA extraction using the AllPrep DNA/RNA Mini kit according to the manufacturer's instructions (QIAGEN). The DNA was then sonicated with the ultrasonicator ME220 (Covaris) and the DNA fragmentation was assessed by 4200 TapeStation (Agilent). Proprietary primers and probes (TaqMan) were designed to target Woodchuck Hepatitis Virus Posttranscriptional Regulatory Element (WPRE) sequence within the CAR construct. The telomerase reverse transcriptase (TERT) gene was used as the reference gene (4458369, Thermo Fisher Scientific). The probes for digital PCR were designed with two different fluorochromes (FAM for WPRE and VIC for TERT) to allow multiplex analysis. Digital PCR assays were performed on the QIAcuity One platform (QIAGEN). 300 ng of DNA/well for a total of 3 wells (of a 24-well QIAcuity Nanoplates 26k), were amplified per sample using the QIAcuity Probe PCR kit (250102, QIAGEN). The conditions of the 2-step dPCR were as follows: 95 °C for 2 min and 35 cycles of 95 °C for 15 s and 60 °C for 30 s. Data was analysed using the QIAcuity Software Suite (QIAGEN). Finally, the CAR-cTreg concentration (copies/µg of DNA) was calculated as described before[9].

**Human CAR screening by Jurkat reporter cell line**

Jurkat (NFAT-Luciferase) reporter cells (60621, Tebubio) were transduced to express different CAR constructs. Here, we used a lentiviral vector backbone containing CD8a hinge, CD3/CD28 costimulatory domain, additional FOXP3 expression cassette and marker gene GFP. They were co-cultured with ENTPD3-expressing cell line RT4 (endogenously expresses ENTPD3) or HEK293T cells engineered to express ENTPD3 at 5:1 and 1:1 ratios respectively. Alternatively, CAR+ Jurkat reporter cells were co-cultured with 2 μg/mL immobilized ENTPD3 peptide (4400-EN, Biotechne). Luciferase activity was measured after 24h with Clariostar Plus (BMG Labtech) luminescence microplate reader.

**CAR activation and suppression assay**

Human Tregs were transduced with CARs or left untranduced (mock) and expanded as described below. Here, we used a lentiviral vector backbone containing CD8a hinge, CD3/CD28 costimulatory domain, additional FOXP3 expression cassette and marker gene GFP. At day 14 a portion of CAR Tregs were stained for anti-G4S (PE, Cell Signalling Technology), CD4 (BV510, OKT-4, Biolegend), CD25 (PE-Cy7, BC96, BD Biosciences), CD39 (BV711, A1, Biolegend), CD62L (PECy594, DREG-56, BD Biosciences), CD73 (BV785, AD2, Biolegend), CTLA4 (BV421, BN13, Biolegend), FOXP3 (AF647, 206D, Biolegend), HELIOS (AF488,22F6, Biolegend), ICOS (BV605, C398.4A, Biolegend) and LAG3 (APC R700, 259D, BD Bioscience). For phenotypic analysis cells were gated on CD4+ and CAR transduced cells. Percentage of indicated markers were assessed. Remaining CAR Tregs were rested for 24 hours and either used for analysis of upregulation of activation markers upon stimulation or suppressive capacity. To check for upregulation of activation markers CAR Tregs were co-cultured with ENTPD3 extracellular domain protein, ENTPD3 expressing HEK293T cells, RT-4, EndoC-βH5 cells (Human Cell Design) or controls; aCD3/aCD28 beads (Dynabeads, Thermo Fisher Scientific), WT HEK293T and no stimulation. Treg activation markers CD69 (PE-Cy7, FN50, Biolegend), CD137 (APC, 4B4-1, Biolegend), and GARP (BV421, 7B11, Biolegend) were assessed by flow cytometry. For analysis of suppressive capacity CAR Tregs and untransduced Mock Tregs were co-cultured with a WT B cell line, ENTPD3 expressing B cell line or aCD3/28 beads (Dynabeads, Thermo Fisher Scientific) and decreasing numbers of CTV-labelled (Thermo Fisher Scientific) CD4 T effector cells. After 5 days cells were assayed by flow cytometry and proliferation was determined by CTV dilution of CD4 Teffs. Percentage Suppression for each ratio of Tregs:Teffs was calculated using the following formula:

$$\text{Suppression} = 100 - (\alpha / \beta * 100)$$

where α is the frequency of proliferating effector T cells in the presence of Treg and β is the frequency of proliferating effector T cells in the absence of Treg cells.

**Generation and stimulation of human CAR Tregs**

Human Tregs were isolated from leukocyte reduction system chambers (LRSC) by pre-enrichment of CD25+ cell population by AutoMACS (Miltenyi) using CD25 MicroBeads II (Miltenyi) followed by cell sorting of CD4+ (FITC, SK3, BD Biosciences) CD25+ (PE, 2A3, BD Biosciences) CD127$^{low}$ (BV421, A019D5, Biolegend) population using FACS Fusion or Aria cell sorters (BD Biosciences). Tregs were incubated and activated in human Treg medium supplemented with 10% human serum and 1% penicillin−streptomycin (Gibco)) supplemented with 1000 U recombinant IL-2 per mL and human Treg expansion beads (Miltenyi) in 1:4 cell/bead ratio for 48 h. Cells were transduced as described above with human CAR vectors and incubated further for 72 h followed by cell sorting of CD4 (BV421, RPA-T4, Biolegend) RQR8 (FITC, QBEND/10, Invitrogen) cell population by FACS Fusion or Aria cell sorters (BD Biosciences). CAR Tregs were either used for TSDR analysis or stimulated further in human Treg medium supplemented with 1000 U recombinant IL-2 per mL in 96-well plates precoated with extracellular domain (Gln 44-Pro 485) of human ENTPD3 protein (Sino Biological).

CAR Tregs activated with human Treg expansion beads (Miltenyi) by 1:4 cell/bead ratio or left unactivated served as positive or negative control, respectively. After 72 h supernatant was saved for cytokine array and cells were used for isolation of mRNA or flow cytometry analysis. For the later CAR Tregs were stained for live cells (Fixable viability dye eFluor 780, Invitrogen), CD4 (BV510, SK3, BD Biosciences), FOXP3 (AF488, PCH101, eBioscience), GARP (APC, 7B11, Biolegend), CD71 (APC, Okt 09, Invitrogen) and CTLA4 (BV421, BNI3, Biolegend) and analysed by flow cytometry. Supernatant of stimulated CAR Tregs was saved for cytokine analysis by LEGENDplex™ Human Essential Immune Response Panel (Biolegend). Samples were 1:5 prediluted and analysed according to the manufacturer's instructions.

**TSDR analysis**

For characterization of demethylation profile of CAR Tregs within Treg specific demethylated regions (TSDR) human nTregs were transduced with ENTPD3-specific h003 or PE-specific Control CAR or left untransduced and cultivated for 3 days with human Treg expansion beads (Miltenyi). gDNA of human CAR Tregs sorted for CAR+ cells were isolated using DNeasy Blood & Tissue Kit (QIAGEN). For bisulfite conversion, the EZ DNA Methylation Gold Kit (Zymo Research) was used by following the manufacture's protocol, with 200ng of gDNA as input. Subsequently, PCR was performed using 12 µl BC converted DNA, 15 µl KAPA HiFi HotStart Uracil+ReadyMix Kit (Roche), 0,9 µl of each primer (10µM, Fw: ACACTCTTTCCCTACACGACGCTCTTCCGATCTTTTGGGGGTAGAGGATTTAGAGGG, Rev: GACTGGAGTTCAGACGTGTGCTCTTCCGATCTCCACATCCACCAACACCCAT) and 1,2 µl of nuclease-free water per sample. Primers are bisulfite-sequence converted and contain Illumina compatible universal adaptor sequences attached at the 5´-end. Thermocycling conditions started with an initial denaturation at 95°C for 3 min, followed by 35 cycles of 1 min each at 98°C, 60°C, and 72°C and ended with a final extension at 72°C for 1 min. Amplicons were purified using the DNA Clean & Concentrator-5 Kit (Zymo Research) according to the manufacturer's protocol. For sequencing, the "Amplicon EZ" service by Azenta Genewiz in Leipzig was used.

**Transcriptome analysis of human ENTPD3-specific CAR Tregs**

mRNA of activated human CAR Tregs was isolated using the RNeasy Mini Kit (QIAGEN) combined with gDNA removal by RNase-Free DNase Set (QIAGEN). Quality and integrity of total RNA was controlled on Agilent Technologies 2100 Bioanalyzer (Agilent Technologies). The RNA sequencing library was generated from 50ng total RNA using NEBNext® UltraTM II Directional RNA Library Prep Kit for Illumina® (New England BioLabs) and NEBNext Poly(A) mRNA magnetic Isolation module according to manufacturer's protocols. The libraries were sequenced on Illumina NovaSeq 6000 using NovaSeq 6000 S1 Reagent Kit (100 cycles, paired end run) with an average of 3 x$10^{7}$ reads per RNA sample. Before alignment to reference genome each sequence in the raw FASTQ files were trimmed on base call quality and sequencing adapter contamination using fastq-mcf. Reads shorter than 15 bp were removed from FASTQ file. Trimmed reads were aligned to the reference genome using open-source short read aligner STAR[10]. Data were compiled using software environment R.

**Human islet microtissue experiments**

For analysis of human ENTPD3-specific CAR interaction with human islets CD8 T cells were transduced with ENTPD3 CARs (h003, h007 and h008). Mock transduced, HLA-A2 CAR transduced and HLA-A2–restricted preproinsulin (PPI)-specific Cytotoxic T lymphocytes (PPI-CTL) CD8 T cells were prepared as controls. GFP was co-expressed with the CAR constructs on all CD8 T cell cohorts (HLA-A2 CAR-GFP fusion protein or co-expressed from the same plasmid with 2A linker, respectively). Prior to co-culture initiation, 3D InSight™ Human Islet microtissues from a single HLA-A2+ donor were stained with Tag-It Violet Proliferation Cell Tracker Dye (425101, Biolegend) for 30 min at 37 °C. CAR T cells and PPI-CTLs were thawed and rested for 2 hours in RPMI 1640 medium at 37 °C and labelled with CellTracker™ Red CMTPX dye (C34552, Thermo Fisher Scientific) for one hour at 37°C with gentle mixing every 20 min. Labelled T cells and hIsMT were washed with medium and then seeded at 2400 cells per hIsMT / per well in Human Islet Maintenance Medium (hIsMM) with 5.5 mM glucose. Co-culture plates were incubated for 72 h in humidified incubator at 37 °C and live imaged.

After 72h co-culture the supernatants were collected for the analysis of IFN-γ and TNF-α secretion. Cytokine secretion was determined according to the protocol provided by the Luminex Performance Human Fixed XL Cytokine Panel 2-Plex kit. The collected co-culture supernatants were diluted 1:2.5, stained for IFN-γ and TNF-α and read within 90 minutes on the Luminex® analyser.

After supernatant collection, hIsMT were washed with Kreb's Ringer Hepes Buffer (KRHB) containing 2.8 mM glucose and then equilibrated in KRHB containing 2.8 mM glucose for 1 hour. Glucose stimulated Insulin Secretion (GSIS) was performed in KRHB containing 16.7 mM glucose for 2 hours and the supernatants were collected. The tissues were then lysed to analyse the total insulin content using Promega CellTiter-Glo® Luminescent Cell Viability Assay with protease inhibitors. Following proper dilutions in KRHB, total and stimulated insulin were be quantified using STELLUX® Chemi Human Insulin ELISA (Alpco).

**Statistical testing**

Statistical significance was determined either by One-way ANOVA or Two-way ANOVA followed by Tukey's multiple comparison test if not stated otherwise. For ANOVA null hypothesis was rejected at $p = 0.05$. If not otherwise stated all replicates are biological replicates corresponding to individual donors. Statistic testing and graphic visualization was performed and rendered by GraphPad 10 (Dotmatics)

**References (Methods)**


1. Fasolino, M., *et al.* Single-cell multi-omics analysis of human pancreatic islets reveals novel cellular states in type 1 diabetes. *Nat Metab* **4**, 284-299 (2022).
2. Pliner, H.A., Shendure, J. & Trapnell, C. Supervised classification enables rapid annotation of cell atlases. *Nat Methods* **16**, 983-986 (2019).
3. Kugler, J., *et al.* Generation and analysis of the improved human HAL9/10 antibody phage display libraries. *BMC Biotechnol* **15**, 10 (2015).
4. Glaser, V., Karsli-Ünal, Ü., Hagedorn, M. & Pieper, T. Antibody Selection on Cells Targeting Membrane Proteins. in *Methods in molecular biology (Clifton, N.J.)*, Vol. 2702 315-325 (2023).
5. Tucker, D.F., *et al.* Isolation of state-dependent monoclonal antibodies against the 12-transmembrane domain glucose transporter 4 using virus-like particles. *Proc Natl Acad Sci U S A* **115**, E4990-E4999 (2018).
6. Saetzler, V., *et al.* Development of Beta-Amyloid-Specific CAR-Tregs for the Treatment of Alzheimer's Disease. *Cells* **12**(2023).
7. Hooijberg, E., Bakker, A.Q., Ruizendaal, J.J. & Spits, H. NFAT-controlled expression of GFP permits visualization and isolation of antigen-stimulated primary human T cells. *Blood* **96**, 459-466 (2000).
8. Pieper, T., *et al.* Generation of Chimeric Antigen Receptors against Tetraspanin 7. *Cells* **12**(2023).
9. Pabst, T., *et al.* Analysis of IL-6 serum levels and CAR T cell-specific digital PCR in the context of cytokine release syndrome. *Exp Hematol* **88**, 7-14 e13 (2020).
10. Dobin, A., *et al.* STAR: ultrafast universal RNA-seq aligner. *Bioinformatics* **29**, 15-21 (2013).

### Supplemental Figures

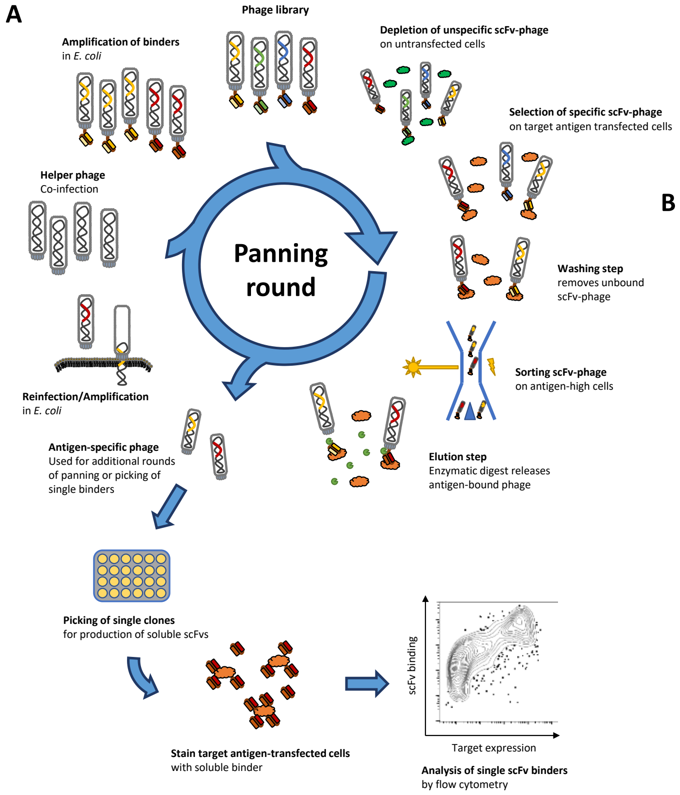


**B**

| | Ectonucleoside triphosphate diphosphohydrolase 3 (ENTPD3) |
|---|---|
| **Family** | E-type nucleotidases (NTPases) |
| **Function** | Ectonucleoside triphosphate diphosphohydrolases which dephosphorylates ATP and adenosine 5'-diphosphat (ADP) |
| **Additional tissue expression** | Brain, spleen, placenta, prostate |
| **Extracellular Domains** | 1 extracellular domain |
| **Total length** | 529 aa (84% extracellular) |
| **Lengths of extracellular domains (ECD)** | 442aa |
| **Homology to murine protein** | 81% (82% ECD) |

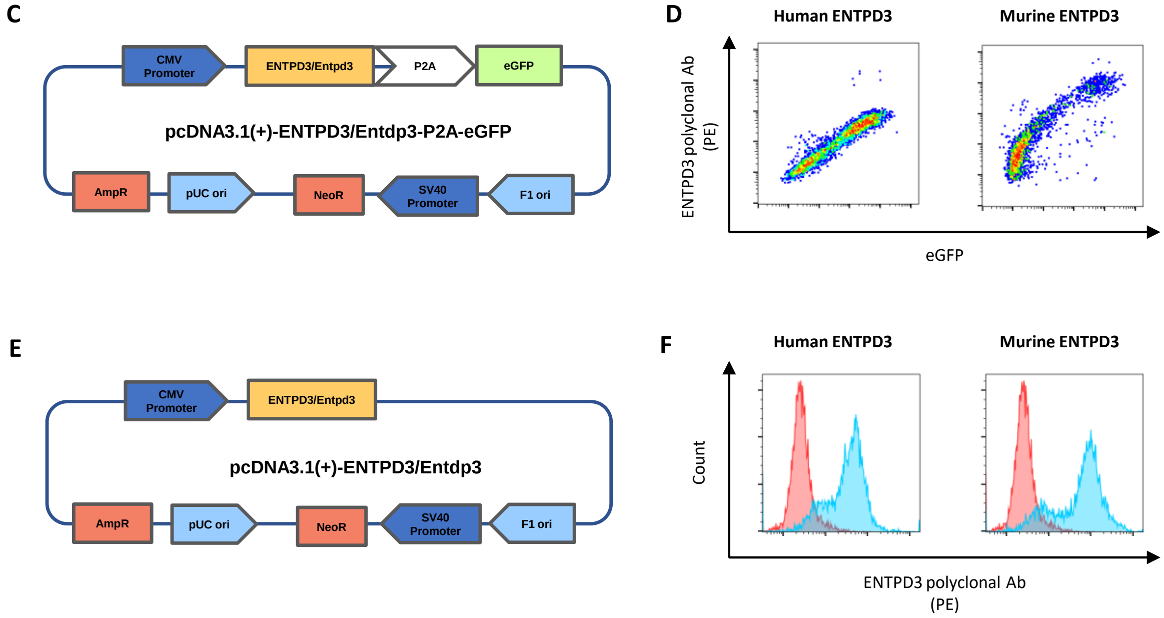


**Supplemental Figure 1:** *Cell-based Phage display as approach for the generation of ENTPD3 specific-scFv binders.* **(A)** Schematic overview of cell-based phage display panning. Human phage library pHAL9/10 was incubated on untransfected HEK293T cells. Unbound scFv-phage were incubated on ENTPD3-expressing cells. Cells were enriched by flow cytometry cell sorting for cells expressing high levels of antigen. Bound scFv-phage were eluted by enzymatic digest and used for amplification in *E. coli* promoted by helper phage coinfection. Panning procedure was repeated for three rounds. In the last round phage-infected single *E.coli* clones were used for the production of soluble scFv. Specificity of scFv was analysed by flow cytometry. **(B)** Functional characteristics and expression of the islet antigen ENTPD3. **(C+E)** Design of target expression vectors. Human ENTPD3 or murine Entpd3 were cloned into pcDNA3.1(+) expression vectors solely (E) or followed by P2A-site separated eGFP (C). **(D)** ENTPD3 targets are expressed on HEK293T cells after transfection with pcDNA3.1(+)-ENTPD3/Entdp3-P2A-eGFP indicated by anti-ENTPD3 antibody staining (PE) and eGFP expression. **(F)** ENTPD3 targets are expressed on HEK293T cells after transfection with pcDNA3.1(+)-ENTPD3/Entdp3 indicated by anti-ENTPD3 antibody staining (PE).

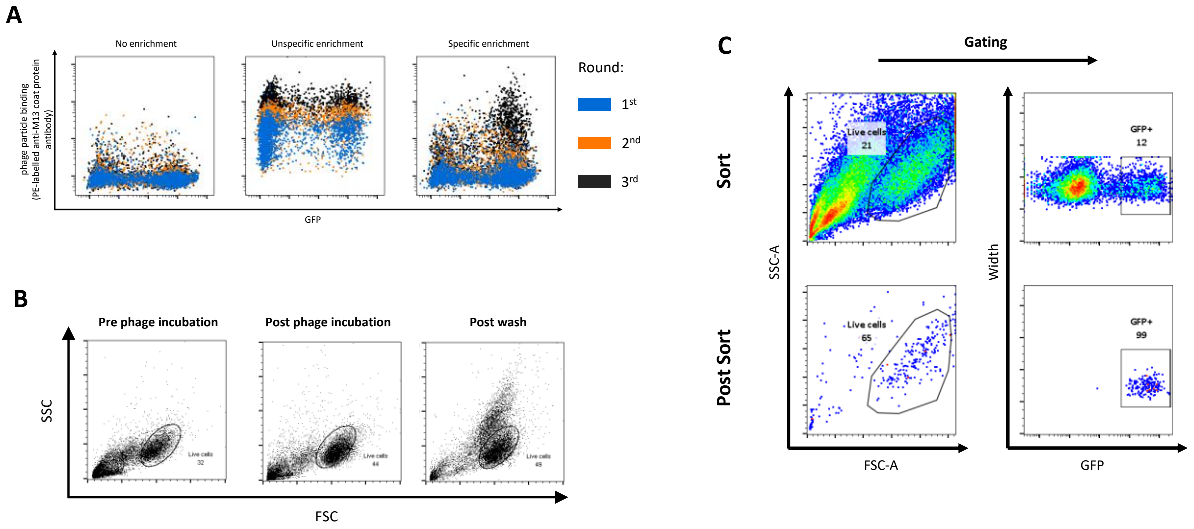


**Supplemental Figure 2:** *Characteristics of panning process* **(A)** Binding of phage pools obtained after consecutive panning rounds. Target:eGFP expressing HEK293T cells were incubated with scFv-phage particles and bound phage were stained with anti-M13 antibody (PE). Results exemplarily shown for a panning approach yielding no, unspecific or specific scFv-phage particles, respectively, of three consecutive rounds. **(B)** Influence of PBS washing buffer (pH5) containing Tween20 on cell viability in the panning process. Cell populations shown in FSC/SSC scatter plot. Note cell population with increased granularity after washing (post sort). **(C)** Sorting strategy for cell panning approach. Cells were sorted for distinct SSC/FSC gate first, followed by sorting of GFP+ population.

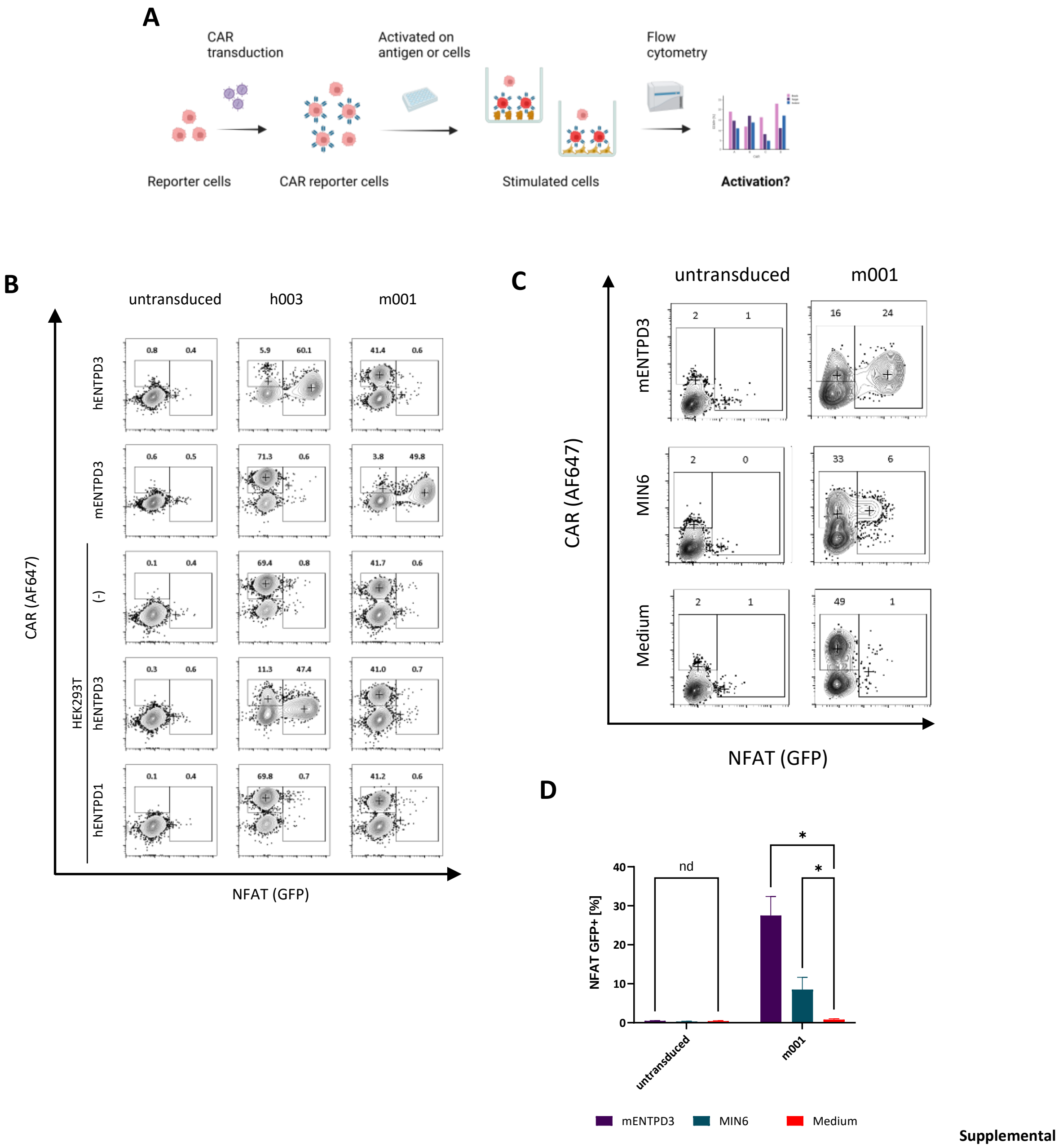


**Supplemental**

**Supplemental Figure 3:** *NFAT:GFP reporter cell assay for CAR screening* **(A)** Schematic overview. **(B)** h003 and m001 CARs analysed in NFAT:GFP reporter cell assay. CAR expression indicated by anti-Fab antibody staining (AF647) and activation (NFAT) indicated by GFP expression after stimulation with human ENTPD3, human ENTPD1, murine ENTPD3 and HEK293T cells transfected for human ENTPD3 or murine ENTPD3 expression, respectively. Untransduced cells served as control. **(C)** Reactivity of murine m001 on murine beta cell line MIN6. Stimulation with murine ENTPD3 protein and medium shown as control. n=3. **(D)** Quantification of C. p values determined by two-way ANOVA and multiple comparison testing (Tukey's test). P values for all experiments: * P < 0.033, ** P < 0.002, *** P < 0.001.

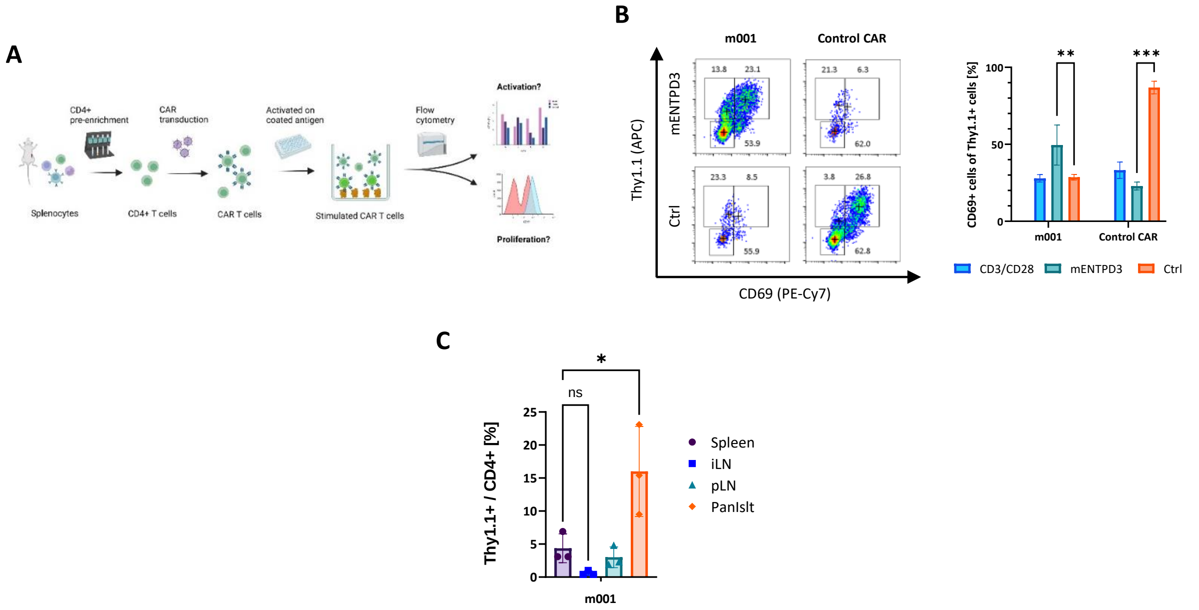


**Supplemental Figure 4:** *Murine m001 CAR Tregs* **(A)** Schematic overview and design of CAR T cell activation assay. **(B)** Activation of CAR T cells on target, aCD3/CD28 beads or control antigen (PE), indicated by expression of CD69 (PE-Cy7). Left: Representative dot plots. Right: Quantification of % CD69+ of Thy1.1+ cells. Control (PE-specific) CAR shown as comparison. Data are presented as mean ± SD, triplicates. p values determined by two-way ANOVA and multiple comparison testing (Tukey's test). **(C)** CAR cTregs enrich specifically in the pancreatic islets (PanIslt) at experimental endpoints. Spleen (Sp), inguinale lymph nodes (iLN) and pancreatic lymph nodes (pLN) shown for comparison. Percentage of Thy1.1+ cells of CD4 cells in respective organs. n = 3 individuals. p values determined by one-way ANOVA and multiple comparison testing (Tukey's test). P values for all experiments * $P < 0.033$, ** $P < 0.002$, *** $P < 0.001$.

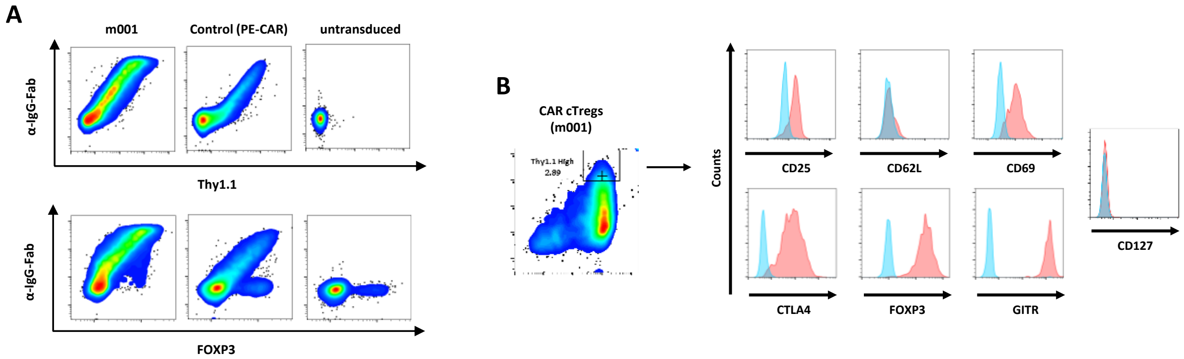


**Supplemental Figure 5:** *Phenotype of murine m001 CAR cTregs* **(A)** Murine m001 CAR cTreg cells of C57Bl/6 were stained for reporter gene Thy1.1, CAR expression (anti-IgG-Fab antibody; upper) and FOXP3 expression (below). **(B)** m001 CAR cTregs of C57Bl/6 were analysed by flow cytometry for expression of T lymphocyte markers. Cells were gated for CD4+ and Thy1.1+. CD25, CD62L, CD69, CD127, CTLA4 (CD152), FOXP3 and GITR expression is shown for high transduced (Thy1.1high) CAR-cTregs in red. FMOs are depicted as internal controls in blue. Plots were normalized to mode.

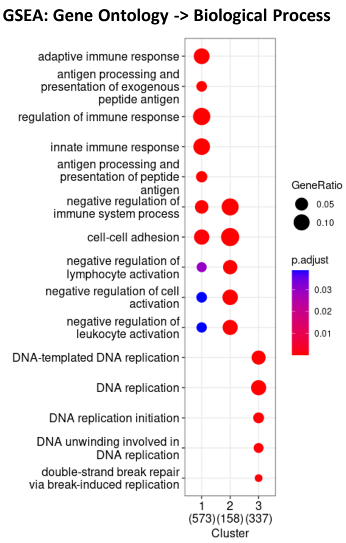


**Supplemental Figure 6:** Gene ontology analysis of major clusters 1, 2 and 3. Size of dots represent ratio of represented genes within each cluster. P value is color coded.

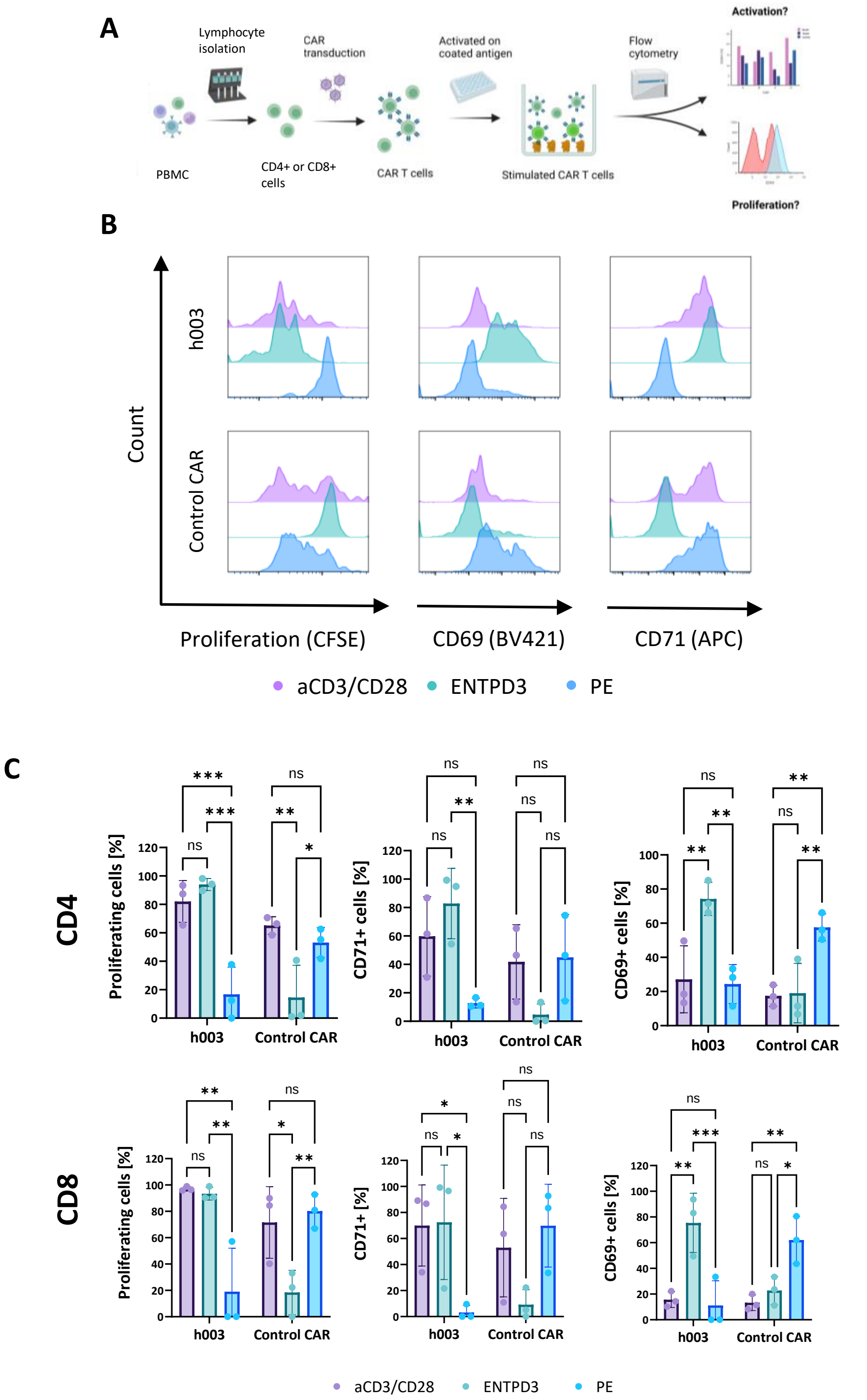


**Supplemental Figure 7: (A)** CD4 and CD8 cells were isolated from PBMC by fluorescence cell sorting and transduced with h003 (ENTPD3) and Control CAR (PE-CAR). CFSE labelled CAR T cells were stimulated on immobilized ENTPD3 , or by aCD3/CD28 bead stimulus. After 96 h cells were analyzed for activation (CD69 and CD71) and proliferation (CFSE) by flow cytometry. **(B)** Proliferation and activation of CD4 and CD8 CAR T cells measured as dilution of CFSE signal or expression of CD69 (BV421) and CD71 (APC), respectively. Cells were cultivated in 3 conditions: Purples: aCD3/CD28 bead stimulus. Green: ENTPD3. Blue: PE. Cells gated on RQR8+ cells. Counts normalized to mode. Representative plots. **(C)** Quantification of (B). Percentages of proliferating (left), CD71+ (middle) and CD69+ (right) CD4+ and CD8+ CAR T cells, respectively. Purples: aCD3/CD28 beads. Green: ENTPD3. Blue: PE. Cells gated on RQR8+ cells. Data presented as mean ± SD of 3 donors. p values determined by two-way ANOVA and multiple comparison testing (Tukey's test). P values for all experiments * $P < 0.033$, ** $P < 0.002$, *** $P < 0.001$.
